# Fluorescent Peptide-based Probe for the Detection of Alpha-synuclein Aggregates in the Gut

**DOI:** 10.1101/2023.11.06.565731

**Authors:** Rachel Sim, Jeremy Lee, Joey Chieng, Ko Hui Tan, Grace Lim, Aaron Foo, Sunny Hei Wong, Kah Leong Lim, Kaicheng Liang

## Abstract

**Background:** Parkinson’s disease (PD) is diagnosed clinically by motor symptoms, with no molecular diagnostic test currently available. By the time motor symptoms manifest, significant irreversible neurodegeneration has already occurred, limiting the effectiveness of neuroprotective therapies and drug interventions. Recent identification of pathological alpha-synuclein (α-syn) aggregates in the gastrointestinal (GI) tract of prodromal PD patients offer a potential avenue for early disease diagnosis. This study aims to explore specific fluorescence labelling of α-syn aggregates in the GI tract using a peptide-based probe for early diagnosis of PD.

**Methods:** We used primary hippocampal neuronal cells and wild-type mouse tissues with the addition of pre-formed α-syn fibrils to identify the most suitable peptide fluorescent probe (**P1**) for staining α-syn aggregates in cells and tissues. We validated the probe labelling in GI tract tissues from three mouse models, including PFF-injected mice and two transgenic PD mouse strains. We quantified labelling accuracy by confocal imaging and protein analysis.

**Results:** We found that **P1** labelled α-syn aggregates with high accuracy (87% in comparison to Serine129-phosphorylated α-syn antibody) and high specificity for labelling their aggregated forms over monomeric forms. In GI tract tissues, **P1** labelled α-syn aggregates across tissue layers (mucosa, sub-mucosa, muscularis externa) and achieved comparable performance to antibody staining. Higher degree of probe labelling was found in older mice due to increased accumulation of α-syn aggregates with ageing. Notably, α-syn aggregates were readily detectable in the colonic mucosae using **P1**, indicating the potential use of this probe for early PD diagnosis during colonic examinations like colonoscopy.

**Conclusion:** We have developed a peptide-based fluorescent probe and demonstrated its rapid and specific labelling of α-syn aggregates. We highlight the probe’s ability to label these aggregates rapidly over α-syn monomers and survey the abundance of α-syn aggregates throughout the entire length of the GI tract. These support the further development of **P1** as a specific fluorescent imaging biomarker for colonic α-syn aggregates for the early detection of PD.

## Background

Parkinson’s disease (PD) is a neurodegenerative disease that is characterised by the loss of dopaminergic neurons in the substantia nigra and presence of intraneuronal Lewy bodies containing aggregated forms of alpha-synuclein (α-syn).[1] PD is currently clinically diagnosed based on the presentation of motor symptoms on a Unified Parkinson’s Disease Rating Scale (UPDRS) and there is no molecular-based diagnostic test available.[2] Current treatments focus on symptom palliation through dopamine replacement therapies and are by no means protective against disease progression.[3] Success in forestalling PD induced neurodegeneration has been largely limited by the severe extent of neuronal loss – approximately 50 to 70% of dopaminergic neurons in the substantia nigra pars compacta are lost by the time the disease is diagnosed.[4, 5] Earlier diagnosis of PD is crucial for increasing the window in which disease-modifying treatments, especially neuroprotective therapies, can be tested. This would improve the success of clinical interventions and aid disease management.

In the prodromal phase, PD patients experience non-motor symptoms such as idiopathic anosmia and gastrointestinal (GI) tract dysfunction.[6] GI manifestations, including dysphagia, vomiting and constipation, can emerge up to two decades before clinical diagnosis of PD.[7] Disruption of GI motility coincides with the emergence of α-syn pathology in the enteric nervous system (ENS) of the GI tract, including the esophagus,[8], stomach,[9] colon[10] and appendix.[11] α-Syn exists physiologically as a soluble monomer but becomes misfolded in pathological states, leading to aggregation and Lewy body formation in cells.[12] Braak and Hawkes have hypothesised that pathological α-syn propagates cranially in a prion-like manner along the vagal axis and affects the brainstem and cerebrum in an ascending fashion as the disease progresses.[13, 14] In keeping with this hypothesis, colonic mucosal biopsy and polypectomy specimens collected in prodromal PD patients demonstrated positive immunostaining for aggregated α-syn 2 to 5 years before the onset of motor symptoms.[15] In 56% of prodromal PD patients, pathological phosphorylated α-syn was also found to have transgressed the mucosal and submucosal layers to seed the intestinal myenteric plexus, which is in contiguity with vagal nerve endings.[16] Further corroborative evidence is availed by comparing the proportion of α-syn inclusion positive gastric biopsy samples in prodromal PD versus advanced PD patients, showing a substantial increase in late-stage disease (17% versus 85% respectively).[17] These findings collectively suggest that pathological aggregated α-syn in the GI tract could be a promising clinical biomarker for the early detection of PD.

Various strategies have been employed to detect α-syn aggregation in PD patients. Immunoassays such as enzyme-linked immunosorbent assay (ELISA) use conformation-specific α-syn antibodies to quantify α-syn aggregates with high sensitivity.[18] In terms of antibody-free approaches, seed amplification assays (SAA) have shown promise for detecting α-syn aggregates in biological fluids such as cerebrospinal fluid[19, 20] and blood.[21] There is also increasing interest in developing fluorescent probes for imaging α-syn fibrils, with recent studies demonstrating utility in living neuroblastoma cells[22] and PD brain sections.[23, 24] However, the utility of fluorescent probes in labelling α-syn aggregates in the GI tract has not been explored.

In this study, we develop a highly specific fluorescence imaging probe that can distinguish between functional monomeric and pathological aggregated forms of α-syn. We validate the binding of the fluorescent probe to different forms of α-syn, and demonstrate its specificity for labelling pathological aggregated α-syn over monomeric α-syn in cellular and animal models. We conduct an in-depth analysis of α-syn labelling in the GI tract, from proximal to distal end, and through the different transmural tissue layers with a focus on quantification of surface mucosal expression. We envision that this peptide-based imaging biomarker can be incorporated into routine colonoscopy screening regimes to identify early-stage PD patients, facilitating more timely clinical interventions to slow the disease progression prior to the development of its classic motor symptoms.

## Methods

### Synthesis of P1

Peptides (GenScript) were modified with a cysteine on the N-terminus. Tris(2-carboxyethyl)phosphine (TCEP) was added to PSMα3 (10 mg/mL in ultrapure water). After 15 min, BODIPY630/650-maleimide in DMSO was added and the mixture was sonicated and left to shake at 1000 rpm at room temperature for 4 hours. The BODIPY630/650-labelled peptide was purified on a desalting column. Fractions containing the labelled peptide were identified using liquid chromatography–mass spectrometry (LC-MS), and the fractions were combined and freeze-dried to give **P1** as a dark blue powder.

### Formation of Alpha-Synuclein (α-syn) Oligomers and Pre-formed Fibrils (PFF)

Mouse α-syn monomers (Proteos, #RP-009) were dialysed against ultrapure water at 4°C overnight using microdialysis tubes (Thermo Fisher Scientific, #69580) with 2-3 buffer replacements to remove the stock buffer solution. Monomers were frozen, lyophilised overnight, and stored in -80°C until further use.

For type A* oligomers generation – The lyophilised α-syn monomers (6.08 mg/mL in PBS) were incubated with ten molar equivalents of (-)-epigallocatechin-3-gallate (EGCG) for 48 h at 37°C, in darkness and without agitation. The solution was then subjected to six consecutive filtration cycles through 100 kDa centrifuge filters (Merck, Darmstadt, Germany) to remove excess compounds and unreacted monomers.

For type B* oligomers generation – The lyophilised α-syn monomers (11.6 mg/mL in PBS) were incubated for 24 h at 37°C without agitation. The solution was subjected to four consecutive cycles of filtration through 100 kDa centrifuge filters. The final concentration of oligomeric solutions was determined by bicinchoninic acid (BCA) assay. The oligomers were kept at -80°C until further use.

For pre-formed fibrils (PFF) generation – The lyophilised α-syn monomers (5 mg/mL in PBS) were incubated in a thermomixer at 1000 rpm, 37°C for 7 days. The solution was subjected to three consecutive cycles of filtration through 100 kDa centrifuge filters. The final concentration of fibrillar samples was determined by measuring the absorbance at 280 nm using a molar extinction coefficient of 7450 M^-1^cm^-1^. Additionally, fibrils were sonicated with SONICS The Vibra-cell VCX-130 for 60 s with the following parameters: 1 s on, 1 s off, 20% amplitude, to generate fibrillar samples with a relatively homogenous size distribution of small fibrils.

### Transmission Electron Microscopy

α-Syn oligomers and PFF samples were placed on glow-discharged grids for 1 minute and grid blotted dry, followed by negative staining with 1% uranyl acetate for 1 minute. The grid was left to dry completely before imaging. Images were acquired using a FEI Tecnai Spirit G2 transmission electron microscope.

### Spectral Measurements of Probe

2 μM of probe was incubated with α-syn monomers, oligomers and PFFs (10 μM) in a black, clear-bottomed 96-well plate (Grenier 655096), with gentle rocking for 30 minutes at RT to allow the probe-α-syn interaction to reach equilibrium. Excitation and emission spectra of the probe was measured on a fluorescence microplate reader (Tecan Infinite M200 Monochromator) at room temperature. The excitation scan was performed with λ_em_ at 735nm, while the emission scan was performed with λ_ex_ at 590nm. Both scans utilised a step size of 1.

### Polyacrylamide gel electrophoresis (PAGE)

Both denaturing SDS-PAGE and non-denaturing Native-PAGE were performed on 2 μg of purified monomers, oligomers and fibrils using 4–15% Mini-PROTEAN TGX Stain-Free Precast Protein Gels (BioRad). SDS was not included in the sample, gel and buffer of the Native-PAGE runs. For Native-PAGE, NativeMark Unstained Protein Standard (ThermoFisher) was used as protein ladder and samples were run for 1 h. For SDS-PAGE, Precision Plus Protein Unstained Standard (BioRad) was used as protein ladder and samples were run for 3 h. No boiling step was included in both cases. To visualise proteins, Coomassie blue protein dye was used as stain.

### Fluorescence Titration of Probe with α-Syn Species

0.5 μM of probe was incubated together with various α-syn species (0 to 10 μM) in a black, clear-bottomed 96-well plate, with gentle rocking for 30 minutes at RT to allow the probe-α-syn interaction to reach equilibrium. Fluorescence values was measured on a fluorescence microplate reader at room temperature with λ_ex_ at 620nm and λ_em_ at 655nm with a step size of 5. The K_D_ binding curve and was obtained with Prism 9.5.1 software (GraphPad Software, San Diego, CA), and data was fitted using a fluorescent ligand binding equation.

### Primary Neuronal Culture

The protocol to harvest primary hippocampal neurons and add PFFs to induce α-syn aggregation was adapted from literature methods.[25, 26] Brains of postnatal mice (P0-P3) were harvested and placed in dissection medium (HBSS with 1 mg/mL sodium pyruvate, 0.1 % glucose, 10 mM HEPES) on ice. The hippocampus was dissected and washed with dissection medium twice. The tissue was incubated with TrypLE solution in a 37°C water bath for 20 min, and a further 5 min at room temperature with the addition of DNase solution. The digested tissue was washed with plating medium (Neurobasal medium with B27 supplement, Glutamax, 50 U/mL penicillin/streptomycin, 10% FBS) twice, then triturated with a P1000 pipette to dissociate the cells. The resulting cell suspension was filtered through a 40-μm nylon mesh cell strainer. The cells were plated on poly-D-lysine-coated well-plates or glass slides for experiments at a density of 52,000 cells/cm^2^. The medium was changed to neuronal medium (Neurobasal medium with B27 supplement, Glutamax, 50 U/mL penicillin/streptomycin) 24 h after plating, and the cells were left to grow for 7 days. PFFs were added to the cells in 80% fresh neuronal medium (20% of medium in each well was retained) and the cells were left to culture for a further 7 days.

### Control Mouse Tissue Preparation

4-month-old BALB/c nude mice were euthanised and organs were harvested. Organs were washed in PBS and fixed in 4% PFA for 15 minutes. Organs were sectioned to 50 μm using a VF-500-0Z microtome (Precisionary Instruments LLC, US).

### PFF-injected Mouse Tissue Preparation

15 µL of 2 mg/mL α-syn PFF (StressMarq SPR-324) were sonicated in an ultrasonic bath sonicator for 2.5h at the following conditions: sweep, 37Hz frequency, 60% power. Water was changed at 30 mins interval to maintain the water temperature at 15°C.

Injection in brain – 10 µg of sonicated PFF (StressMarq) was injected into the left striatum of 4-6 months old C57BL6 mice using the following stereotaxic coordinates ML +2.0 mm, AP +0.5mm, DV - 3.5mm. 5 µL of 2 mg/mL of sonicated PFF was injected at speed of 1 µL/min. 1-month post-injection, the mice were transcardially perfused with PBS followed by 4% PFA in PBS. The brain was harvested and kept in 4% PFA in PBS overnight. The brain was then kept in 15% sucrose in PBS for 24 hours, followed by 30% sucrose in PBS for another 24 hours, before embedding in optimal cutting temperature (OCT) medium. Cryosectioning was performed at 30 µm thickness.

Injection in gut – 9-11 weeks old C57BL6 mice were injected with 6 µg of sonicated PFF (Proteos) at a total of 4 sites: 2 at the gastric pylorus and 2 at the upper duodenum (2 mg/mL, vol = 3 µL), totaling 24 µg of PFF. 2.5 months later, mice were transcardially perfused with PBS followed by 4% PFA in PBS. The stomach and upper duodenum were extracted and flushed with PBS to remove waste material. Tissues were kept in 4% PFA in PBS overnight, followed by 70% ethanol overnight. The stomach and upper duodenum were separately embedded in paraffin blocks. Paraffin sectioning was performed at 5 µm thickness.

### Transgenic Mouse Tissue Preparation

Two strains of transgenic mice were imported from the Jackson Laboratory: (i) M83 transgenic mouse JAX004479[27] and (ii) double transgenic mouse JAX010799[28]. JAX004479 mice were sacrificed at 10 months old. JAX010799 mice were sacrificed at 5 and 15 months old. The tissues were harvested in this order from sacrificed mice: brain, esophagus, stomach, small intestine (duodenum, jejunum, and ileum) and colon. Intestinal tissues were flushed with PBS multiple times to remove waste material. The intestinal tissues were coiled in a swiss-roll manner[29] on sponges in tissue embedding cassettes (Uni-Sci, C33118001PH) and fixed directly in the cassettes. Stomach was cut open and laid out flat on the sponge in the tissue cassette. Tissues were fixed in 4% PFA in PBS overnight at 4°C, then embedded in paraffin blocks.

### Immunofluorescence (IF) Staining

For cells – Cells were treated with **P1** (1 µM) for 5 min at 37°C, before washing in PBS. The cells were fixed with 4% PFA at room temperature. Immunofluorescence staining steps were performed as follows: Blocking with TBS-T with 2% horse serum for 1 h, incubation with primary antibodies overnight at 4°C, washing with TBS-T (3 times), incubation with secondary antibodies at room temperature for 2 h. The cells were then incubated with Hoechst 33342 for 15 min, washed with TBS and fixed with anti-fade mounting medium (Abcam, #ab104135).

For tissues – Fresh tissues were embedded in 2.5% agarose and sliced at 50 μm thickness using a vibratome. The slices were mounted onto a poly-lysine coated glass slide, air dried and used for subsequent staining. Fixed tissues embedded in paraffin blocks were sectioned onto glass slides. The tissue sections were deparaffinised and rehydrated with sequential dipping into the following solutions: xylene (2 times), 100% ethanol (2 times), 95% ethanol, 70% ethanol, 50% ethanol, distilled water. Antigen retrieval was carried out by incubating the slides in citrate buffer for 1 h at 60°C. Immunofluorescence staining was performed as described for cells. **P1** (1 µM) was added together with Hoechst 33342 (0.01 mg/mL) after the immunofluorescence staining and incubated for 30 min at room temperature. The tissues were washed with TBS and fixed with anti-fade mounting medium.

**Table 1.**
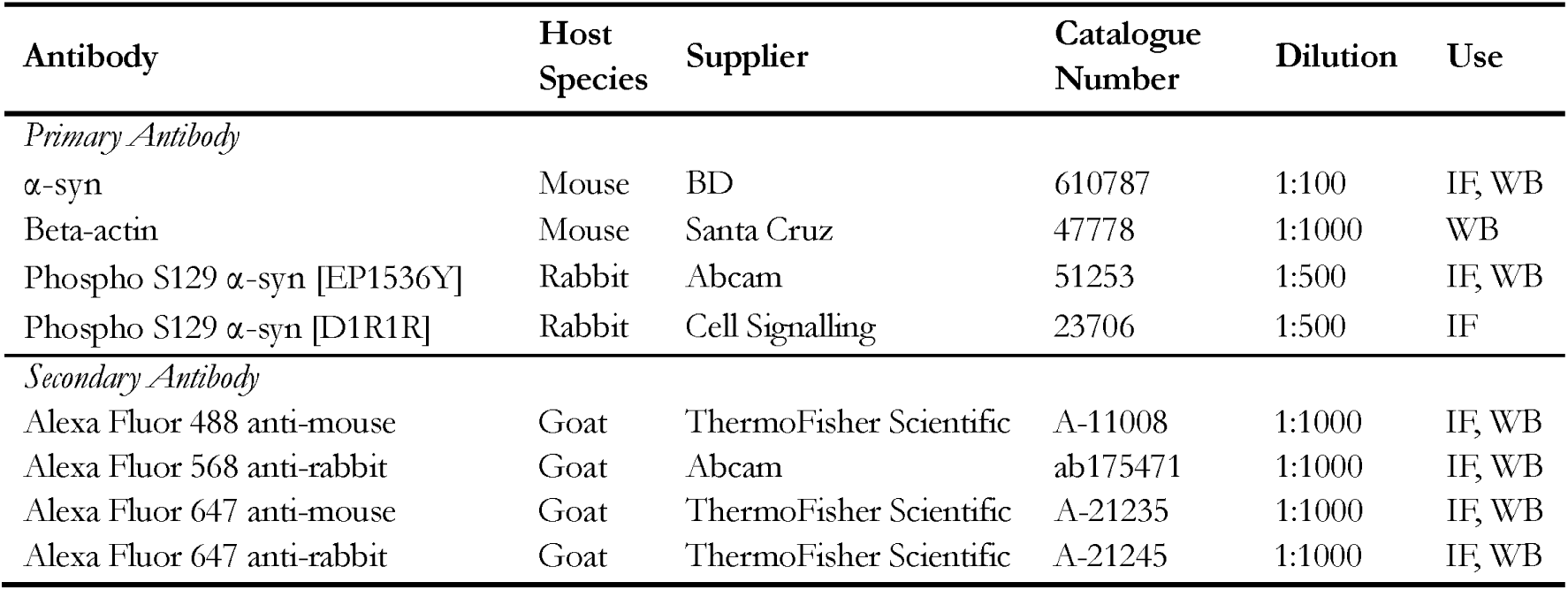
Antibodies used in the study.

| Antibody | Host Species | Supplier | Catalogue Number | Dilution | Use |
| --- | --- | --- | --- | --- | --- |
| *Primary Antibody* |  |  |  |  |  |
| α-syn | Mouse | BD | 610787 | 1:100 | IF, WB |
| Beta-actin | Mouse | Santa Cruz | 47778 | 1:1000 | WB |
| Phospho S129 α-syn [EP1536Y] | Rabbit | Abcam | 51253 | 1:500 | IF, WB |
| Phospho S129 α-syn [D1R1R] | Rabbit | Cell Signalling | 23706 | 1:500 | IF |
| *Secondary Antibody* |  |  |  |  |  |
| Alexa Fluor 488 anti-mouse | Goat | ThermoFisher Scientific | A-11008 | 1:1000 | IF, WB |
| Alexa Fluor 568 anti-rabbit | Goat | Abcam | ab175471 | 1:1000 | IF, WB |
| Alexa Fluor 647 anti-mouse | Goat | ThermoFisher Scientific | A-21235 | 1:1000 | IF, WB |
| Alexa Fluor 647 anti-rabbit | Goat | ThermoFisher Scientific | A-21245 | 1:1000 | IF, WB |

### Microscopy

Fluorescence images were acquired on an Olympus inverted fluorescence IX83 microscope and an Olympus FV3000RS confocal microscope. High content imaging was done on a PerkinElmer Operetta CLS imaging system. Brightfield images were acquired with a Zeiss AxioObserver 7 Microscope.

### Colocalisation Quantification

The Manders colocalisation, Costes colocalisation and Pearson’s correlation coefficients between pS129 staining (orange channel) and P1 staining (red channel) were determined using CellProfiler (v 4.2.1). The images were converted to 8-bit grayscale images and illumination correction was applied. The channel images were aligned, then the correlation between the two channels were measured. The Manders colocalisation and Costes colocalisation coefficients were calculated based on methods described in literature.[30] The Pearson’s correlation measurement is the normalised covariance, which is the covariance divided by the product of standard deviation of pixels in each image. Individual objects were automatically detected, with thresholding size set at a minimum of 3, and a maximum of 15 pixel units. Objects were counted in both channels, and objects found to touch and overlap each other were considered as colocalised. The probe positive predictive value is calculated by the number of objects colocalised in the orange and red channel (true positives), divided by the total number of objects detected in the red channel (probe positives).

### Protein Extraction and Western Blotting

About 10 mg of tissue was flash frozen in dry ice. The frozen tissue was minced and homogenised with a mortar and pestle chilled on ice. Homogenized tissue was suspended in lysis buffer (20 mM Tris-HCl, pH 7.4, 150 mM NaCl, 1% Triton X-100, 2% SDS, 1x Pierce protease and phosphatase inhibitor cocktail). The mixture was sonicated for 5 s for 3 times, 1 s on, 1 s off, 30% amplitude, at 4°C. The samples were centrifuged at 12000 x g at 20°C for 10 min and the supernatant was collected. Concentration of protein sample collected was determined using the Bradford assay. LDS sample buffer and reducing agent was added to 50 μg of protein lysate, and the samples were heated at 70°C for 10 min. The proteins were separated on a 4 to 12% Bis-Tris gel using MES as running buffer, at 200 V for 30 min. The proteins were transferred onto a nitrocellulose membrane on an iBlot 2 gel transfer (dry blotting) system, at 20 V for 1 min, 23 V for 4 min and 25 V for 2 min. The membrane was fixed with 4% PFA for 30 min, then washed with TBS and blocked with TBS-T containing 2% horse serum for 1 h. The membrane was incubated with primary antibodies overnight at 4°C and washed with TBS-T (3 times). The membrane was then incubated with fluorescent-labelled secondary antibodies for 2 h at room temperature. The membrane was imaged on an iBright imaging system.

### Statistical Analysis

All values are expressed as the meanlJ±lJS.E.M. Differences between means were evaluated by two-way ANOVA test (Prism, Graph Pad Software, San Diego, CA).

## Results

### BODIPY630/650 as the fluorophore for optimized labelling

We synthesised a specific peptide-based fluorescent imaging marker to label pathological α-syn aggregates. (Fig 1a) Cationic α-helical peptide PSMα3 was previously reported to bind toxic α-syn oligomers and amyloid fibrils with low nanomolar affinity, over the monomeric functional α-syn protein.[31] We first chose Boron Dipyrromethene (BODIPY-FL) as the fluorescent tag for functionalization of PSMα3, due to its neutral charge, low molecular weight, and high quantum yield in conjugates. BODIPY-FL has been used for *in vivo* imaging applications[32] and has been incorporated into a fluorescent probe undergoing clinical trial for detection of oral cancers.[33] While BODIPY-FL-labelled **P0** showed specific labelling of α-syn aggregates in cells, (Fig S1a) BODIPY-FL fluoresces in the green channel that is prone to tissue auto-fluorescence, resulting in high background in tissue staining. (Fig S1b) To avoid background from tissue auto-fluorescence, we explored using far-red dyes with longer excitation and emission wavelengths. The cyanine5.5 dye resulted in non-specific labelling contributing to high background signal and was deemed unsuitable for use. (Fig S1c) We eventually chose BODIPY630/650 as the fluorescent tag as it is photostable and demonstrates minimal non-specific background labelling. BODIPY630/650-labelled **P1** was obtained after gel filtration chromatography with high yield of 90%. (Fig S2a) A single peak corresponding to the expected mass of **P1** in the liquid chromatography-mass spectrometry (LC-MS) spectrum confirms that the peptide is labelled with BODIPY630/650. (Fig S2b) As the reaction is quick, single-step and high-yielding, the synthesis of **P1** can be rapidly scaled up for larger quantities needed in pre-clinical or clinical stages. The fluorescent and physiochemical properties of the probe can also be further fine-tuned if required, by modifications in the chemical structure of the BODIPY group.

**Figure 1.**
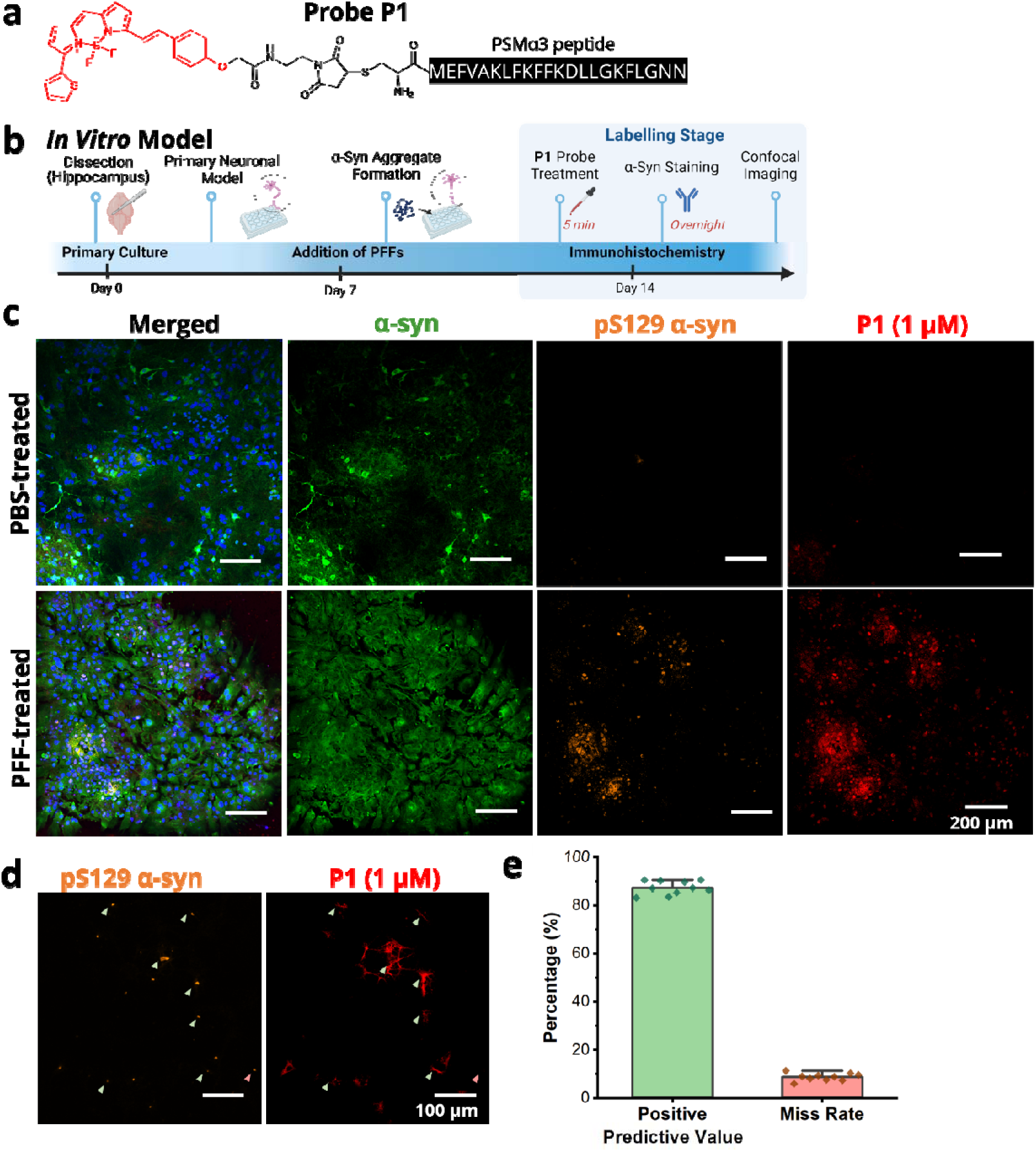
Probe P1 shows high labelling accuracy for α-syn aggregates in vitro, compared to phosphorylated (pS129) α-syn antibody which was used as the gold standard reference staining for α-syn aggregates. (a) Structure of peptide-based probe **P1**; (b) Setup of primary hippocampal neuronal culture and induction of α-syn aggregation upon addition of PFFs; (c) Confocal microscopy of primary neuronal cells post 7 days of PFF addition. Nuclei in blue, total α-syn in green, pS129 α-syn in orange, and probe **P1** in red. Scale bar represents 200 μm; (d) Probe **P1** shows colocalised staining with the PS129 α-syn antibody staining. The probe labels most of the same focal locations, but with slightly larger surrounding area than the PS129 α-syn antibody hits – this is still regarded as a positive colocalisation. True positives (indicated by green arrows) are stained by both **P1** and pS129 α-syn antibody. False negatives (indicated by pink arrows) are not stained by **P1** but stained by pS129 α-syn antibody. Scale bar represents 100 μm; (e) Graph showing average positive predictive value (true positives/probe positives) of 87.4% and miss rate (false negatives/antibody positives) of 8.7%. Values on graph are given as the mean.LJ±.LJSEM calculated over 10 wells, with 9 fields of view each.

### P1 shows high labelling specificity for α-syn aggregates induced in primary hippocampal neurons

We determined the labelling specificity of fluorescent probe **P1** in an *in vitro* model established using primary hippocampal neurons prepared from postnatal mice pups to closely mimic a neuronal environment (Fig 1b). α-Syn preformed fibrils (PFFs) were generated from mouse recombinant α-syn[25] and used to induce aggregate formation in primary neurons. The addition of PFFs causes endogenous α-syn in primary neurons to assemble into Lewy bodies-like aggregates that are hyperphosphorylated and have a filamentous ultrastructure. Primary neurons inoculated with PFFs for 7 days were subsequently treated with **P1** for 5 min, for the labelling of α-syn aggregates, before fixation and immunofluorescence staining. An antibody specific for phosphorylation at serine-129 (pS129) in α-syn aggregates [Abcam EP1536Y] was used as a gold-standard to visualize α-syn aggregates and an α-syn antibody [BD610787] was used to visualise total α-syn including monomeric and aggregated α-syn. In control cells that were treated with PBS, α-syn monomers were present (green), while there was no significant staining by **P1** (red), indicating that the probe did not label α-syn monomers. In cells that were treated with PFFs, presence of aggregated α-syn was confirmed by pS129 α-syn antibody staining. (Fig 1c) There was good colocalisation between pS129 α-syn antibody staining (orange) and **P1** staining (red), indicating that **P1** probe labelled α-syn aggregates with high specificity over functional monomeric α-syn. (Fig 1d) The positive predictive value (PPV) of the probe was determined by the number of true hits (labelled by both the probe and pS129 α-syn antibody, highlighted by green arrows in Fig 1d) over the number of positive hits (labelled by the probe). The average PPV was calculated to be 87.4%. The miss rate was determined by the number of false negatives (aggregates not labelled by the probe but labelled by the pS129 α-syn antibody, highlighted by pink arrows in Fig 1d) over the number of positive hits. **P1** had a low miss rate of 8.7%. (Fig 1e)

### P1 shows preferential binding to B* oligomers and fibrils over A* oligomers and monomers

Prior work has demonstrated the ability of PSMα3 to bind toxic α-syn aggregates, including both oligomeric and fibrillar forms.[31] To investigate if the incorporation of the BODIPY630/650 moiety in **P1** affected the binding affinity of PSMα3 to toxic α-syn aggregates, we characterised the interaction between **P1** and various α-syn species. Various α-syn aggregates (non-toxic A* oligomers, toxic B* oligomers, PFFs) were prepared from mouse α-syn monomers according to literature[34-36] and their size and morphology were characterised by polyacrylamide gel electrophoresis (PAGE) and transmission electron microscopy (TEM). (Fig S3a,b)

We studied the general fluorescence properties of **P1** by measuring the excitation and emission spectra of **P1** in the presence of α-syn monomers, oligomers and fibrils. (Fig 2a,b) The excitation peak of **P1** increased by 1.8-fold and 7.6-fold in intensity upon binding to the B* oligomers and PFFs respectively. Similarly, its emission peak also increased by 3-fold and 16-fold in intensity upon binding to the B* oligomers and PFFs respectively. Fluorescence enhancement upon probe binding to the α-syn aggregates indicates that the probe can specifically detect aggregates over monomeric species. This phenomenon is most likely due to the PSMα3 peptide binding to the hydrophobic β-sheet regions of B* oligomers and fibrils.[31] This places the hydrophobic BODIPY moiety within the hydrophobic microenvironment of the β-sheets, where they form non-covalent interactions. These interactions result in restricted rotation of the fluorophore and shield the fluorophore from non-radiative decay of fluorescence from water molecules in the aqueous media,[37] contributing to the increased fluorescence observed.

**Figure 2.**
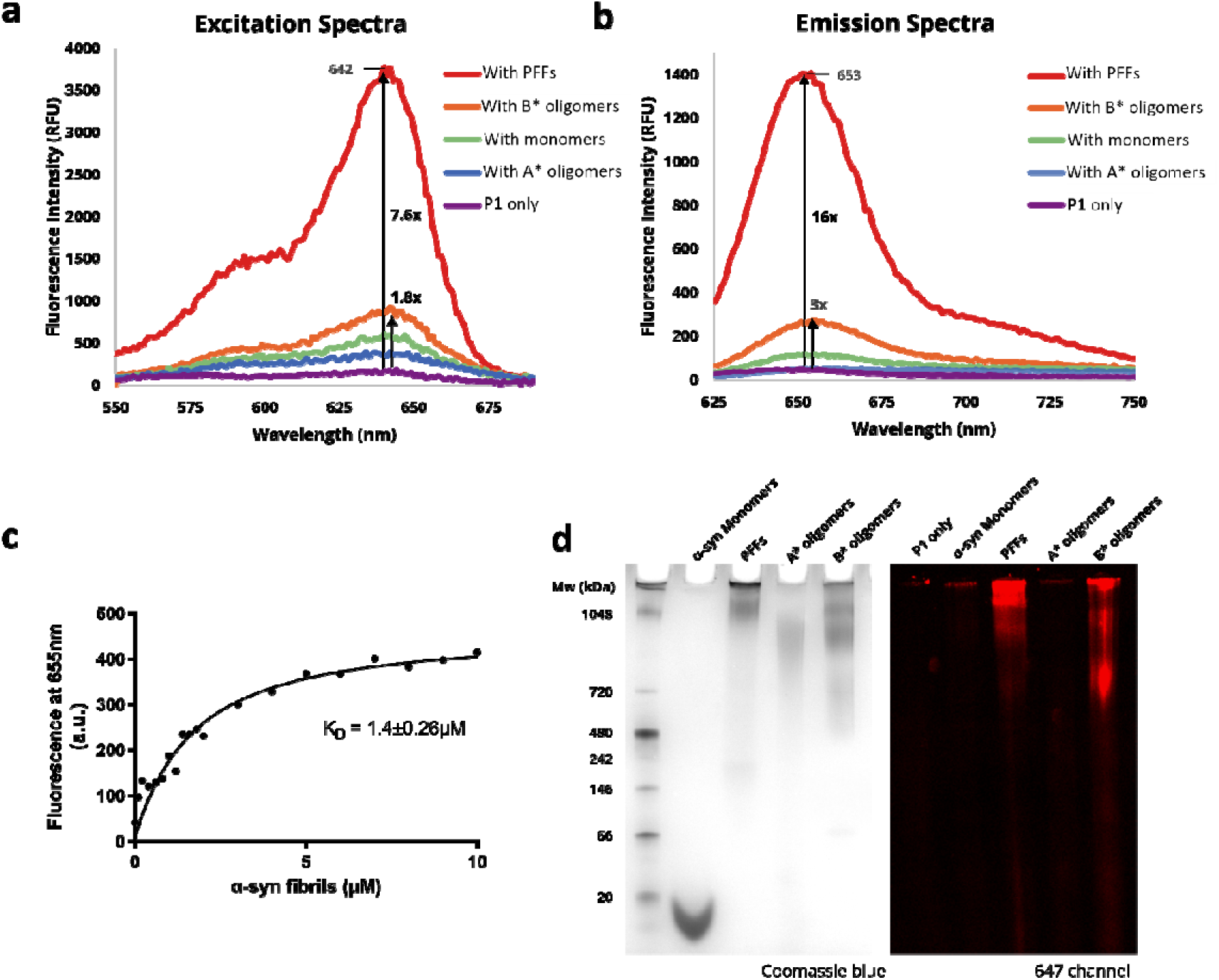
Probe P1 exhibits preferential binding to α-syn fibrils over α-syn monomers. (a) Fluorescent excitation spectra and (b) fluorescent emission spectra of **P1**. 2 µM of probe **P1** was incubated with 10 µM of various α-syn species, including monomers, A* oligomers, B* oligomers and PFFs. Concentrations of all α-syn species are expressed at its equivalent monomer concentration. Fluorescence intensity of **P1** increases significantly upon interaction with PFFs (7.6x and 16x for excitation and emission respectively); (c) Fluorescence titration curve of **P1** (0.5 µM) to PFFs, showing micromolar K_D_; (d) Native-PAGE images of **P1** with various α-syn species, showing that **P1** preferentially binds to B* oligomers and PFFs, over α-syn monomers and A* oligomers. 2 µg of **P1** was incubated with 2 µg of α-syn monomers and various α-syn aggregate species and run under native PAGE conditions. In-gel fluorescence was first visualised in the 647 channel, then the gel was subsequently imaged in the brightfield channel after staining with Coomassie blue. Red bands correspond to **P1** bound to the B* oligomers and PFFs.

We performed fluorescence titration by titrating **P1** against a range of concentrations of the various α-syn species. (Fig 2c, Fig S4) **P1** was found to bind to PFFs with a K_D_ of 1.4±0.26 µM. This K_D_ value is higher than the reported nanomolar binding affinity of PSMα3 for fibrils, determined directly by highly sensitive spectroscopic techniques, Förster Resonance Energy Transfer (FRET) and fluorescence cross-correlation spectroscopy (FCSS).[31] In our fluorescence titration assay, we relied on indirect measurements of BODIPY fluorescence gain when the probe is bound to the ligand.[38] Insufficient fluorescence enhancement was observed at nanomolar concentrations of the probe, hence we were unable to determine K_D_ values below the micromolar range. Nevertheless, we demonstrated that **P1** exhibited the strongest binding affinity to α-syn fibrils over other α-syn species.

We visualised probe-α-syn complexes using Native-PAGE to preserve the native conformation of the aggregates, as we found that the higher molecular weight aggregates were partly dissociated upon separation on SDS-PAGE (Fig S3a). In-gel fluorescence was observed corresponding to the B* oligomers and PFFs, indicating that **P1** interacts specifically with the toxic aggregate forms. (Fig 2d) The α-syn monomers and non-toxic A* oligomers showed no fluorescence, indicating that **P1** did not bind to these non-toxic α-syn species. This shows that **P1** exhibits higher binding specificity for toxic α-syn aggregates containing β-sheet regions.

### P1 detects α-syn aggregates in PFF-injected brain and gut tissues with low background signal

We next sought to evaluate the use of **P1** in *ex vivo* staining using tissues obtained from PFF-injected wild-type mouse models. (Fig 3a) PFFs were injected directly into the brain of wild-type mouse to induce the formation of α-syn aggregates. A high degree of α-syn aggregation was observed in the hippocampal and cortex regions, labelled both by pS129 antibody and **P1**. (Fig 3b) No significant α-syn aggregation was observed in the control group, where the mice were injected with PBS. (Fig S5) High background signal was observed in the green channel, which is attributed to tissue autofluorescence. When BODIPY-FL-labelled **P0** which fluoresces in the green channel was tested, we found that it was very challenging to distinguish between the probe staining and the tissue autofluorescence. (Fig S1b) This highlights the importance of using a far-red dye to minimise background signal and avoid false positive results.

**Figure 3.**
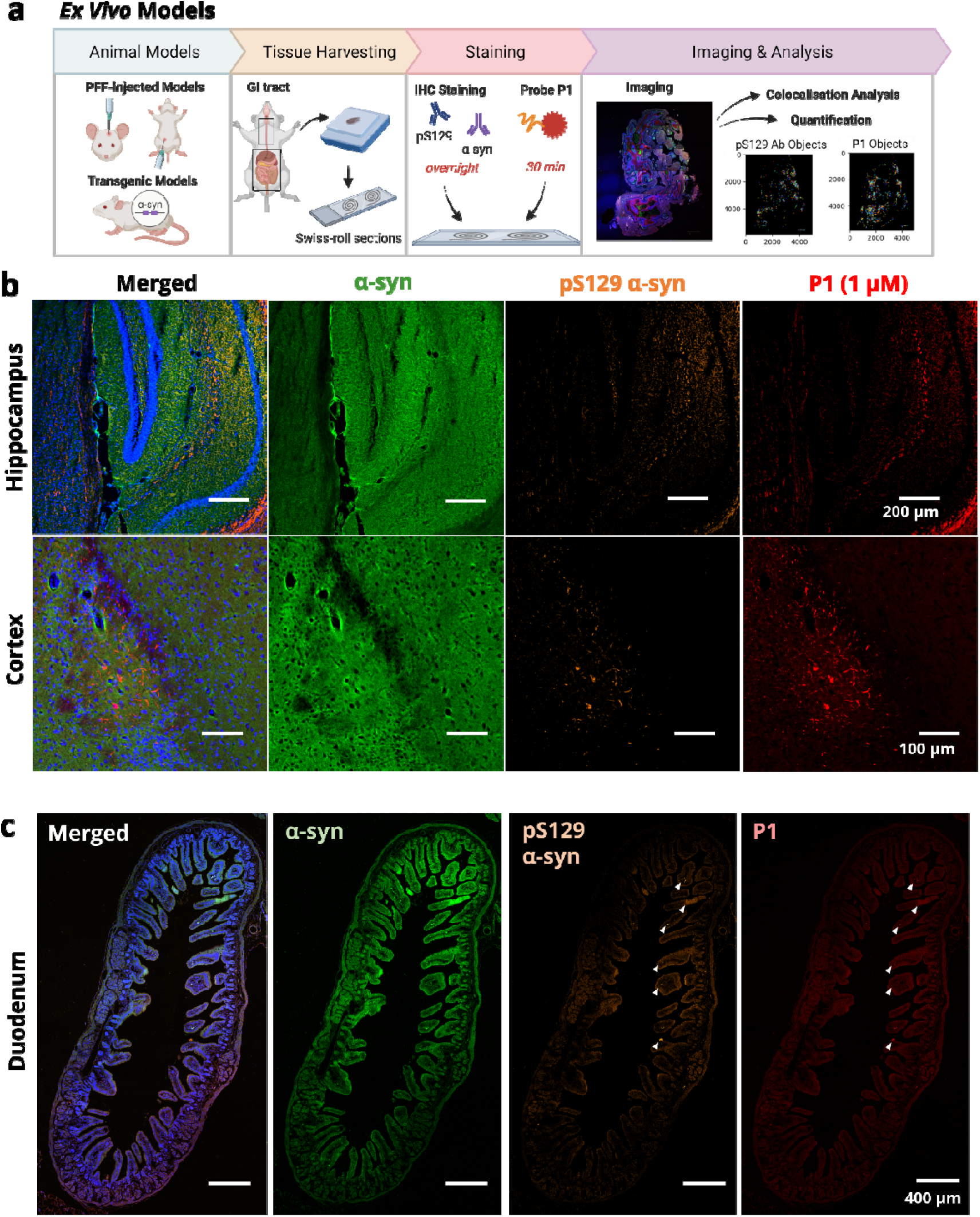
Probe P1 shows specific staining of α-syn aggregates in tissues from PFF-injected mouse models. (a) Models used for ex vivo tissue labelling: PFF-injected mouse, where PFF was injected directly into the brain or duodenum, and transgenic mouse models overexpressing human α-syn; (b) Confocal microscopy of brain tissues post 30 days of PFF injection showing high levels of α-syn aggregation in the hippocampal and cortex regions, labelled specifically by both pS129 α-syn antibody and **P1**; (c) Confocal microscopy of duodenum tissues post 75 days of PFF injection showing low levels of α-syn aggregation in the mucosa region indicated by the white arrows. Nuclei in blue, α-syn in green, pS129 α-syn in orange and probe in red. Images of tissues from control PBS-injected mice are shown in Fig S5 for comparison.

We evaluated the *ex vivo* tissue staining of P1 in gut-injected mouse models. Expression of α-syn is observed in the gut of wild-type mouse (Fig S6). It is hence expected that injection of PFFs directly in the gut of wild-type mouse would induce the aggregation of endogenous α-syn. However, only a low degree of α-syn aggregation was observed in the duodenum tissue from gut-injected mouse. (Fig 3c) This could be due to a limitation of the PFF-injected model, as the α-syn aggregation is predominantly concentrated at the site of injection and takes some time to propagate through the enteric nervous system. While some α-syn aggregates could be seen in the mucosa layer, labelled by both pS129 antibody and **P1**, this effect could vary based on the site of PFF injection and hence may not be representative of the distribution of α-syn aggregation across tissue layers in the GI tract.

### P1 highlights differential accumulation of α-syn aggregates across GI tract in transgenic mice

To evaluate *ex vivo* labelling in the GI tract tissues and study the distribution of α-syn aggregation within the tissue layers, we employed transgenic PD mouse models. We selected two transgenic models demonstrating α-syn aggregation: (i) M83 transgenic mouse JAX004479 and (ii) double transgenic mouse JAX010799. JAX004479 mice were reported to have significant α-syn aggregates in the brain and onset of motor phenotype by 8-10 months of age.[27] JAX010799 mouse were reported to have early gastrointestinal dysfunction, before the onset of motor symptoms.[28] To evaluate probe staining, the esophagus, stomach, duodenum, jejunum, ileum and colon tissues from JAX004479 mice were harvested for tissue section staining. By using a swiss-roll preparation and longitudinal sectioning for the harvested intestinal tissues, we could capture a large area of intestinal tissue and different portions from the proximal to the distal end for comprehensive histological assessment. α-Syn aggregation was observed across all the GI tract tissues investigated, with the highest intensity of staining observed in the esophagus and colon. (Fig 4a, b) α-Syn aggregates were found in the mucosa layers, stained by both pS129 and **P1** in a similar pattern. α-Syn aggregates were more prevalent in the extreme proximal and distal ends of the GI tract, with **P1** staining accounting for 12.9% and 13.3% of the total tissue area in the esophagus and colon respectively. Analysis using the Costes method revealed an average of 91.0% colocalisation between the two channels (pS129 and **P1**), indicating that **P1** staining was largely comparable to pS129 antibody staining. (Fig 4c) Using CellProfiler software, objects labelled by both pS129 antibody and **P1** were automatically counted and used to determine the probe positive predictive value (PPV). The PPV was generally consistent across all the GI tract tissues, with an average of 87.8%. (Fig 4d) This demonstrates that the labelling specificity of **P1** was not affected by differences in the environments of the various GI tract tissues. The PPV in tissues was similar to the PPV in cells, indicating that the probe was able to label α-syn aggregates in both cells and tissues with similar sensitivity.

**Figure 4.**
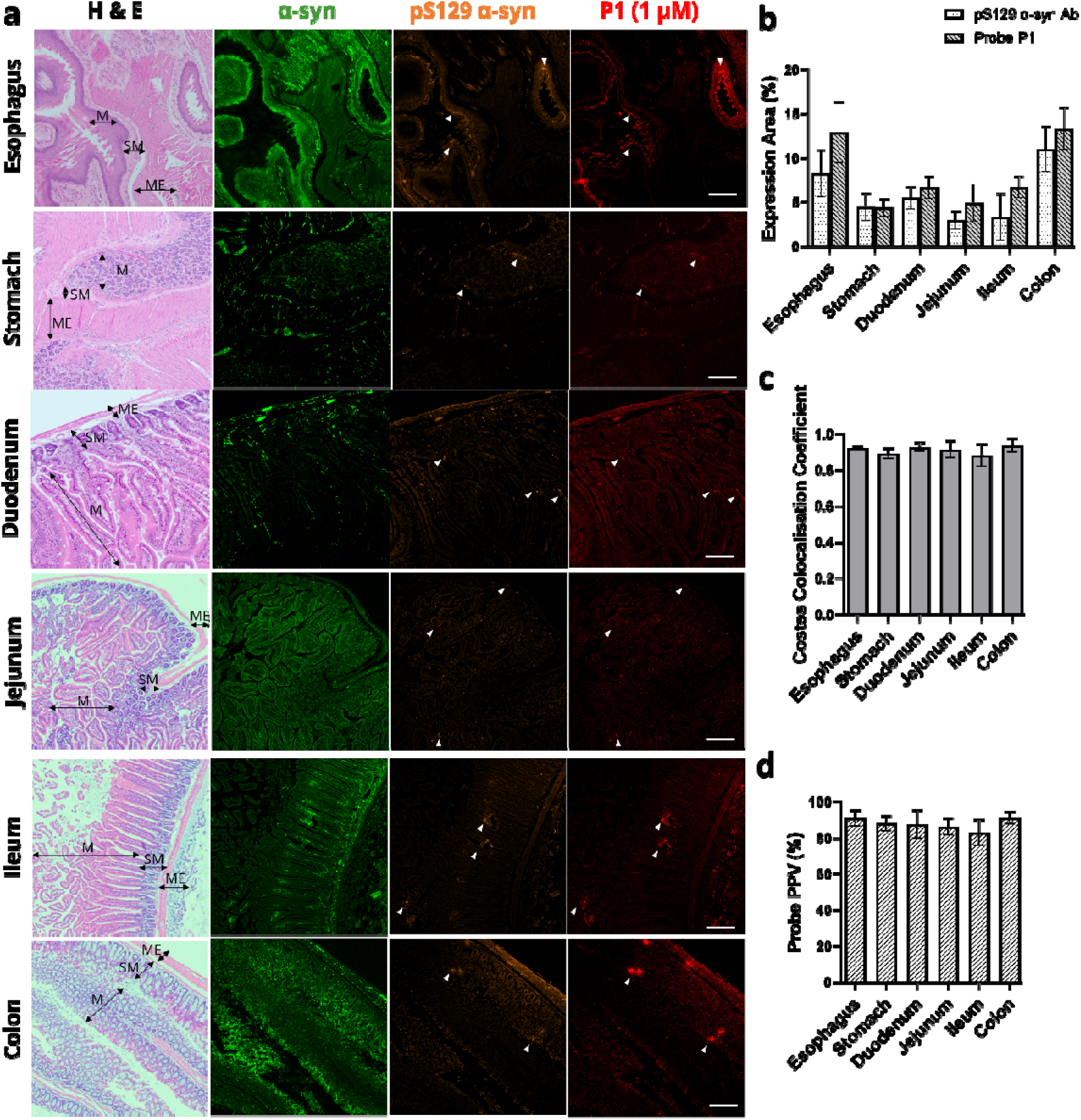
Specific α-syn antibody and probe P1 labelling of entire GI tract (esophagus, stomach, duodenum, jejunum, ileum and colon) in transgenic mice. (a) Aggregated α-syn is observed in the mucosa layers and stained by both pS129 α-syn antibody and **P1** (annotated with white arrows). Tissue layers as indicated by black arrows: muscularis externa (ME), submucosa (SM), mucosa (M). Nuclei in blue, α-syn in green, pS129 α-syn in orange and probe in red. Scale bar represents 200 μm; (b) Total expression areas of α-syn aggregates labelled by pS129 α-syn antibody and **P1** were quantified, with the highest expressions observed in the esophagus and colon; (c) High degree of colocalisation (average of 0.91) was observed between pS129 α-syn antibody and probe **P1** channels across all the GI tract tissues, determined by the Costes method; (d) Probe positive predictive values were consistent across the respective GI tract tissues. Values on graphs are given as the mean.LJ±.LJSEM (n.LJ=.LJ3 per group).

### Accumulation of a-syn aggregation increases with age in transgenic mice

We evaluated the accumulation of α-syn aggregation within GI tract tissues by comparing young and old JAX010799 transgenic mice. The ages of the transgenic mice were chosen based on the presence of motor symptoms: 5-month-old mice presented no motor symptoms, corresponding to early stage PD, while 15-month old mice presented motor symptoms such as tremors, corresponding to late stage PD. Increased levels of total α-syn were observed in the brain and GI tissues of older transgenic mice. Higher molecular weight species of α-syn were also present in increased amounts in the older mice (highlighted by red arrows). There was generally lower abundance of α-syn in lysates obtained from duodenum and colon tissues, which may be due to difficulties in solubilising α-syn aggregates in tissues, or high levels of protein degradation in fresh GI tract tissues during protein extraction protocols. A 2-fold increase in total α-syn in the brain and 1.5-fold increase in the esophagus of 15-month-old mice indicated α-syn accumulation with age. (Fig 5b)

**Figure 5.**
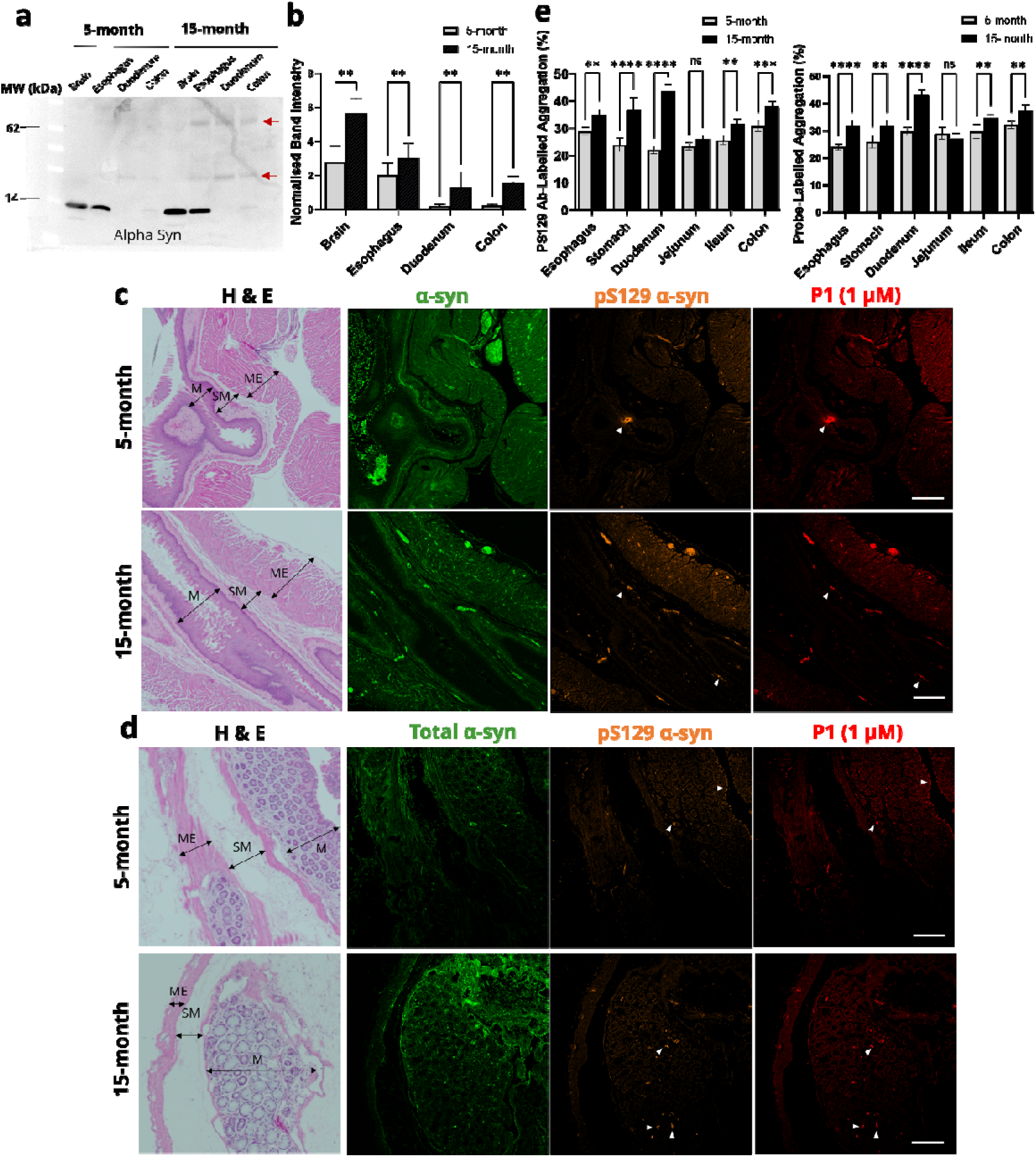
Increased accumulation of α-syn aggregates across GI tissues in older transgenic mice. (a) Western blot showing abundance of total α-syn in brain, esophagus, duodenum and colon of young (5-month) and old (15-month) transgenic mice, with higher molecular weight α-syn species observed in the older mice (indicated by red arrows); (b) Graph showing intensity of bands corresponding to α-syn, normalised to beta-actin, quantified from blots in Fig S7. Older mice showed increased intensities of α-syn bands; Confocal images showing increased staining of aggregated α-syn in (c) esophagus and (d) colon of older transgenic mice. Tissue layers as indicated by black arrows: muscularis externa (ME), submucosa (SM), mucosa (M). Scale bar represents 200 μm; (e) Graphs showing degree of α-syn aggregation determined by pS129 α-syn antibody labelling and **P1** labelling, with generally higher degrees of α-syn aggregation observed across all GI tissues in older mice. Values on graphs are given as the mean.LJ±.LJSEM (n.LJ=.LJ3 per group). Ns: not significant; * p.LJ<.LJ0.05; ** p.LJ<.LJ0.01; *** p.LJ<.LJ0.001; **** p.LJ<.LJ0.0001, determined by two-way ANOVA test.

Comparison of α-syn aggregation between young and old mice was also done with immunohistochemical staining and image analysis. Increased abundance of α-syn aggregates was observed in esophagus and colon tissues of older mice. (Fig 5c,d) The degree of α-syn aggregation was quantified by assessing the ratio of probe **P1** staining to total α-syn antibody staining, and the ratio of pS129 α-syn antibody staining to total α-syn antibody staining. Our findings consistently revealed a higher degree of α-syn aggregation across all GI tract tissues in older transgenic mice. (Fig 5e) The trend was similar in both pS129 α-syn antibody-labelled and probe **P1**-labelled aggregation, demonstrating that our peptide probe shows potential as a diagnostic marker comparable to pS129 α-syn antibody.

### α-Syn aggregates are detectable in colonic mucosa

As endoscopic imaging is limited largely to the mucosal layers,[39] we sought to determine the abundance of α-syn aggregation in colonic mucosa to determine the feasibility of endoscopic detection. α-Syn aggregates were abundant in the mucosa and labelled by both pS129 α-syn antibody and **P1**. (Fig 6a) The highest degree of α-syn aggregation was observed in the muscularis externa compared to the other tissue layers. (Fig 6b) Notwithstanding this, > 20% of α-syn present in the mucosal layer was found to be aggregated and labelled by pS129 α-syn antibody and **P1**. While α-syn aggregates were present in the lowest levels in the mucosa compared to the submucosa and muscularis externa regions, they were still detectable by probe staining. The presence of α-syn aggregates in mucosa indicates that α-syn aggregation is a potential biomarker for endoscopy screening methods.

**Figure 6.**
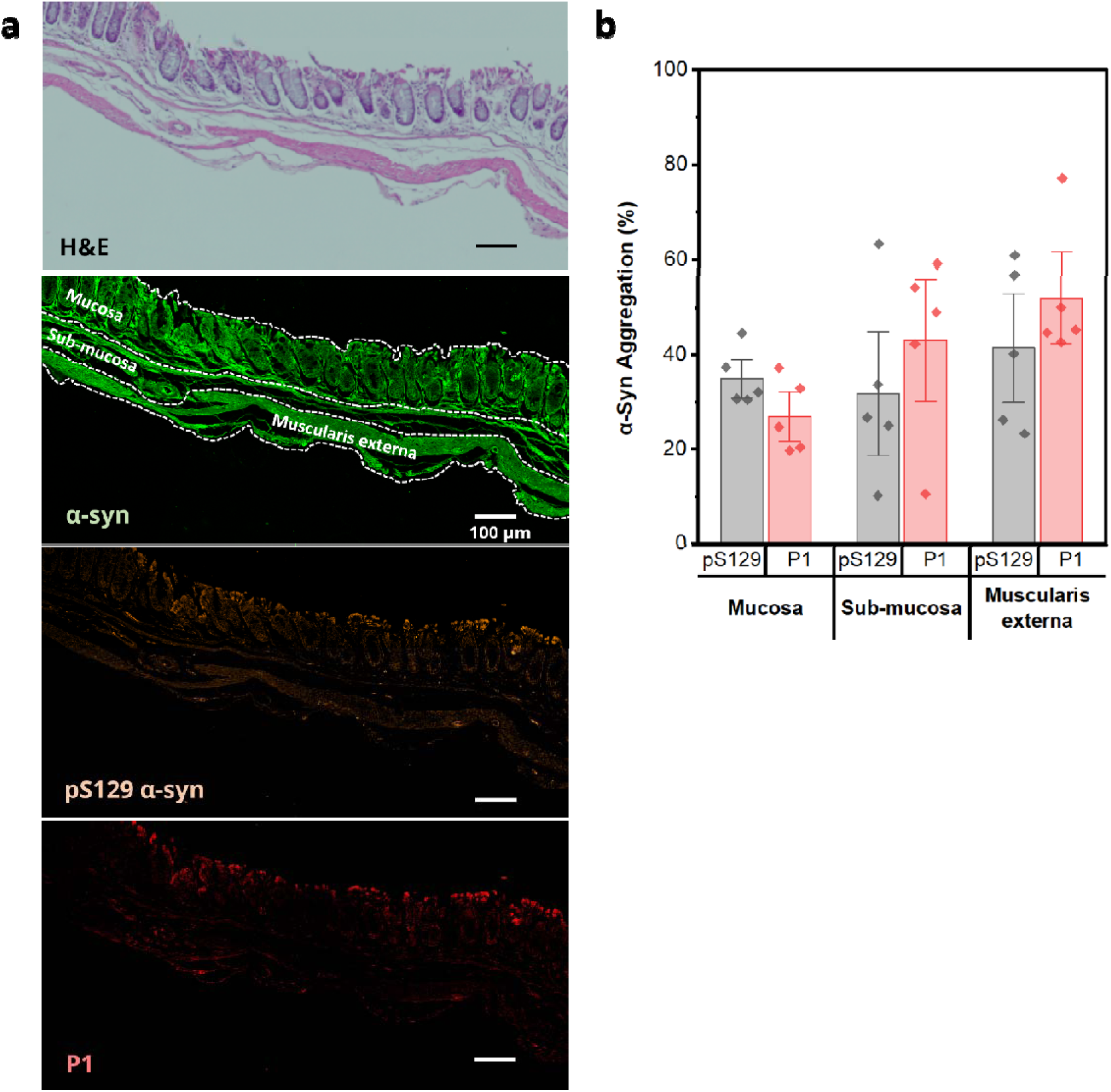
Longitudinal view of unrolled colon tissue of transgenic mouse showing differing levels of α-syn aggregation across tissue layers (mucosa, sub-mucosa and muscularis externa). (a) Confocal microscopy of selected section of colon tissue from transgenic mouse, with the different tissue layers marked by white dashed lines. α-Syn aggregates on the mucosa surface are specifically labelled by pS129 α-syn antibody (orange) and **P1** (red). Full view of section is shown in Fig S8; (b) Graph showing degree of α-syn aggregation (pS129 α-syn antibody or **P1** positives, divided by α-syn positives) in the mucosa, sub-mucosa and muscularis externa regions. Mucosa shows lower but detectable levels of **P1** staining compared to the other tissue layers.

## Discussion

In this study, we demonstrate the use of a peptide-based fluorescent imaging marker to label α-syn aggregates with comparable specificity and sensitivity to an antibody specific for phosphorylated α-syn, in primary neurons, wild-type mouse and transgenic mouse tissue samples. A range of cellular and tissue models were chosen, with consideration that α-syn aggregates are made up of a heterogenous mix of oligomeric and fibrillar species, and that propagation and accumulation of α-syn aggregates differ across models. By using PFFs to induce α-syn aggregation in primary cells and wild-type mouse models, we observed specific labelling of aggregated α-syn over functional monomeric α-syn by probe **P1**. In brain-injected mice, some spread of α-syn aggregation was observed, with α-syn aggregates found in the cortex, away from the striatum which was the site of injection. However, there was limited spread of α-syn aggregation in the gut-injected mice, possibly due to the type of PFFs used or a short time frame of 2.5 months before tissue harvesting. These PFF-injected models have some limitations as the recruitment of endogenous mouse α-syn by PFFs to form higher molecular weight aggregates occurs slowly and is largely dependent on the seeding technique and site of injection.[40]

To demonstrate the utility of the probe in disease models where α-syn aggregation is more sparsely distributed over a larger region, we used transgenic PD mice which overexpress α-syn. As the overexpression of α-syn in the transgenic models is not limited to the GI tract, these mice may not entirely recapitulate the gut-brain pathway for α-syn propagation in PD. However, these transgenic mice display GI dysfunction before central nervous system dysfunction, and are still a relevant model for studying prodromal PD.[41] We demonstrated high *ex vivo* labelling sensitivity in GI tract tissues of transgenic mouse, highlighting the potential of using fluorescence imaging in the GI tract as a detection method for diagnosis. The relatively higher prevalence of α-syn aggregation in the esophagus and colon suggests that gastroscopy and colonoscopy could be leveraged to access these GI tract regions for detection. We observed specific probe staining for α-syn aggregates in GI tract tissues of young transgenic mice which were not exhibiting any obvious motor deficits. This suggests that the probe can be used for detection of α-syn aggregates in the GI tract as an early indication of PD before symptom onset. Comparing GI tract images from young and old transgenic mice, we observed an increase in α-syn aggregation in older mice, which coincides with the manifestation of motor symptoms. This indicates that there is a decline in ability to clear α-syn aggregates from the cells with increasing age, and the higher degree of aggregation may be linked to motor deficits. The increased aggregation is in line with clinical studies, which found that the ratio of pS129 α-syn to total α-syn was higher in cerebrospinal fluid samples with PD progression, and may be used as a diagnostic marker for PD.[42, 43] Furthermore, our findings suggest the possibility of establishing quantification thresholds for probe staining at various stages of PD progression. Such thresholds could transform this imaging biomarker into a staging tool for assessing disease severity in clinical samples. However, this requires further validation through comparisons of probe labelling percentages in the GI tract between healthy controls and PD patients at various disease stages.

Our peptide-based fluorescent probe offers several advantages over the pS129 α-syn antibody, especially for further development as an imaging marker for diagnostic use. Firstly, it is low-cost and much cheaper to produce on a larger scale as compared to antibodies. Secondly, it can be used for live cell and tissue imaging as the peptide can be taken up into cells without the need for fixation and permeabilisation. Thirdly, our probe stains rapidly, and demonstrates increased emission signal upon binding to the target aggregated α-syn species, allowing for the strong fluorescence labelling to be captured quickly by imaging. While SAAs have reported high sensitivities and specificities for distinguishing PD patients from control patients using cerebrospinal fluid samples, these assays are currently primarily qualitative in nature.[44] Challenges persist in obtaining quantitative data due to inherent variability in aggregation processes and kinetics in different assay conditions, making it hard to establish standardised thresholds that correlate with disease severity and progression.[45] In comparison, probe **P1** labels the α-syn aggregates directly in the tissue without relying on amplification processes, potentially allowing for quantification thresholds to be determined and correlated with clinical phenotypes.

While our current study demonstrates the utility of our probe in mouse models, further work to validate the probe staining in human tissue samples will aid clinical translation. The ability to visualise α-syn aggregates within the mucosal layers of GI tract tissues is promising for the development of an endoscopic detection method. A survey of a late middle-aged population with a mean age of 60.3 years found that 11% of the subjects had more than two PD risk factors, and 9.7% had constipation.[46] Our probe could serve as a surveillance tool to screen patients with non-motor symptoms and deemed to be at risk of prodromal PD. The at-risk age for prodromal PD coincides with the recommended age for colorectal cancer screening (50-75 years),[47] which presents an opportunity for the integration of PD screening into routine colonoscopy protocols without any additional inconvenience to patients. A recent study revealed that in intestinal biopsies of PD patients, the ratios of aggregated α-syn to total α-syn in the sigmoid mucosa were notably higher compared to healthy controls.[48] However, the depth of field in endoscopic imaging may be limited to a more superficial mucosa layer, hence *in vivo* studies will provide more insight on the ability to detect α-syn aggregates via endoscopy. More studies will also be required to differentiate the α-syn aggregation patterns among other synucleinopathies such as Lewy body dementia and multiple system atrophy, allowing for the distinction of PD from these related diseases.

## Conclusion

We report a peptide-based fluorescent probe as a specific imaging marker to label α-syn aggregates in cellular and animal models of synucleinopathies. Probe **P1** demonstrates a high positive predictive value of 88%, in comparison to antibodies staining for phosphorylated, aggregated α-syn, in GI tract tissues. Being low-cost, rapid staining and highly specific, **P1** shows promise as a diagnostic tool to evaluate PD risk in GI tissue biopsies or routine endoscopic screening procedures. With further clinical validation, we envision that **P1** can provide a means for PD surveillance in the older population and for classification of PD patients according to disease stages.

## Abbreviations

α-syn: Alpha-synuclein
BCA: Bicinchoninic acid
BSA: Bovine serum albumin
ELISA: Enzyme-linked immunosorbent assay
ENS: Enteric nervous system
FBS: Fetal bovine serum
FRET: Förster Resonance Energy Transfer
FCSS: Fluorescence cross-correlation spectroscopy
GI: Gastrointestinal
H&E: Hematoxylin and eosin
IF: Immunofluorescence
LC-MS: Liquid chromatography–mass spectrometry
min: minutes
PAGE: Polyacrylamide gel electrophoresis
PBS: Phosphate buffered saline
PD: Parkinson’s disease
PFF: Pre-formed fibrils
PPV: Positive predictive value
s: seconds
SAA: Seed amplification assay
pS129: Phosphorylated α-syn at residue serine 129
SDS: Sodium dodecyl sulfate
TEM: Transmission electron microscopy
TBS: Tris buffered saline
TBS-T: Tris buffered saline with 0.1% tween-20
WB: Western blot

## Declarations

### Ethics approval and consent to participate

Animal work was conducted with IACUC approval (#221718).

### Consent for publication

Not applicable

### Availability of data and materials

Additional data are included in the supplementary information file. Raw imaging data acquired during this work are available from the corresponding author upon reasonable request.

### Authors’ contributions

RS led the conceptualisation, experimental work and writing of manuscript; JL, JC, KHT, GL contributed to experimental work, acquisition of data and writing; AF, SHW and KLL contributed to manuscript editing; KL led project direction, supervision, and manuscript editing.

## Acknowledgements

Figures 1b and 3a were created on Biorender.com and exported under licence. The authors thank the Electron Microscopy Unit (https://medicine.nus.edu.sg/core-facilities/electron-microscopy-unit-emu/) in National University of Singapore and the team, particularly Micky Leong, for their support and assistance in the EM work. The authors would like to thank the A*STAR Microscopy Platform (AMP) and the AMP staff for the training, consultation, advice, and technical support. The authors thank the A*STAR Biological Resource Centre (BRC) and their staff for the training and technical support for animal work.

## References

1. Poewe W, Seppi K, Tanner CM, Halliday GM, Brundin P, Volkmann J, Schrag A-E, Lang AE: Parkinson disease. Nature Reviews Disease Primers 2017, 3(1):17013.

2. Armstrong MJ, Okun MS: Diagnosis and Treatment of Parkinson Disease: A Review. JAMA 2020, 323(6):548–560.

3. Salamon A, Zádori D, Szpisjak L, Klivényi P, Vécsei L: Neuroprotection in Parkinson’s disease: facts and hopes. Journal of Neural Transmission 2020, 127(5):821–829.

4. Ross GW, Petrovitch H, Abbott RD, Nelson J, Markesbery W, Davis D, Hardman J, Launer L, Masaki K, Tanner CM et al: Parkinsonian signs and substantia nigra neuron density in decendents elders without PD. Ann Neurol 2004, 56(4):532–539.

5. Dauer W, Przedborski S: Parkinson’s Disease: Mechanisms and Models. Neuron 2003, 39(6):889–909.

6. Chaudhuri KR, Schapira AHV: Non-motor symptoms of Parkinson’s disease: dopaminergic pathophysiology and treatment. The Lancet Neurology 2009, 8(5):464–474.

7. Warnecke T, Schäfer KH, Claus I, Del Tredici K, Jost WH: Gastrointestinal involvement in Parkinson’s disease: pathophysiology, diagnosis, and management. npj Parkinson’s Disease 2022, 8(1):31.

8. Tanei ZI, Saito Y, Ito S, Matsubara T, Motoda A, Yamazaki M, Sakashita Y, Kawakami I, Ikemura M, Tanaka S et al: Lewy pathology of the esophagus correlates with the progression of Lewy body disease: a Japanese cohort study of autopsy cases. Acta Neuropathol 2021, 141(1):25–37.

9. Del Tredici K, Duda JE: Peripheral Lewy body pathology in Parkinson’s disease and incidental Lewy body disease: four cases. J Neurol Sci 2011, 310(1-2):100–106.

10. Beach TG, Adler CH, Sue LI, Vedders L, Lue L, White Iii CL, Akiyama H, Caviness JN, Shill HA, Sabbagh MN et al: Multi-organ distribution of phosphorylated alpha-synuclein histopathology in subjects with Lewy body disorders. Acta Neuropathol 2010, 119(6):689–702.

11. Gray MT, Munoz DG, Gray DA, Schlossmacher MG, Woulfe JM: Alpha-synuclein in the appendiceal mucosa of neurologically intact subjects. Mov Disord 2014, 29(8):991–998.

12. Breydo L, Wu JW, Uversky VN: Α-synuclein misfolding and Parkinson’s disease. Biochim Biophys Acta 2012, 1822(2):261–285.

13. Braak H, Rüb U, Gai WP, Del Tredici K: Idiopathic Parkinson’s disease: possible routes by which vulnerable neuronal types may be subject to neuroinvasion by an unknown pathogen. J Neural Transm (Vienna) 2003, 110(5):517–536.

14. Hawkes CH, Del Tredici K, Braak H: Parkinson’s disease: a dual-hit hypothesis. Neuropathol Appl Neurobiol 2007, 33(6):599–614.

15. Shannon KM, Keshavarzian A, Dodiya HB, Jakate S, Kordower JH: Is alpha-synuclein in the colon a biomarker for premotor Parkinson’s Disease? Evidence from 3 cases. Movement Disorders 2012, 27(6):716-719.

16. Stokholm MG, Danielsen EH, Hamilton-Dutoit SJ, Borghammer P: Pathological α-synuclein in gastrointestinal tissues from prodromal Parkinson disease patients. Annals of Neurology 2016, 79(6):940–949.

17. Sánchez-Ferro Á, Rábano A, Catalán MJ, Rodríguez-Valcárcel FC, Díez SF, Herreros-Rodríguez J, García-Cobos E, Álvarez-Santullano MM, López-Manzanares L, Mosqueira AJ et al: In vivo gastric detection of α-synuclein inclusions in Parkinson’s disease. Movement Disorders 2015, 30(4):517–524.

18. El-Agnaf OM, Salem SA, Paleologou KE, Curran MD, Gibson MJ, Court JA, Schlossmacher MG, Allsop D: Detection of oligomeric forms of alpha-synuclein protein in human plasma as a potential biomarker for Parkinson’s disease. Faseb j 2006, 20(3):419–425.

19. Fernandes Gomes B, Farris CM, Ma Y, Concha-Marambio L, Lebovitz R, Nellgård B, Dalla K, Constantinescu J, Constantinescu R, Gobom J et al: α-Synuclein seed amplification assay as a diagnostic tool for parkinsonian disorders. Parkinsonism Relat Disord 2023:105807.

20. Kang UJ, Boehme AK, Fairfoul G, Shahnawaz M, Ma TC, Hutten SJ, Green A, Soto C: Comparative study of cerebrospinal fluid α-synuclein seeding aggregation assays for diagnosis of Parkinson’s disease. Mov Disord 2019, 34(4):536–544.

21. Kluge A, Bunk J, Schaeffer E, Drobny A, Xiang W, Knacke H, Bub S, Lückstädt W, Arnold P, Lucius R et al: Detection of neuron-derived pathological α-synuclein in blood. Brain 2022, 145(9):3058–3071.

22. Gaur P, Galkin M, Kurochka A, Ghosh S, Yushchenko DA, Shvadchak VV: Fluorescent Probe for Selective Imaging of α-Synuclein Fibrils in Living Cells. ACS Chemical Neuroscience 2021, 12(8):1293–1298.

23. Watanabe H, Ono M, Ariyoshi T, Katayanagi R, Saji H: Novel Benzothiazole Derivatives as Fluorescent Probes for Detection of β-Amyloid and α-Synuclein Aggregates. ACS Chemical Neuroscience 2017, 8(8):1656–1662.

24. Zeng Q, Cui M: Current Progress in the Development of Probes for Targeting α-Synuclein Aggregates. ACS Chemical Neuroscience 2022, 13(5):552–571.

25. Volpicelli-Daley LA, Luk KC, Lee VM: Addition of exogenous alpha-synuclein preformed fibrils to primary neuronal cultures to seed recruitment of endogenous alpha-synuclein to Lewy body and Lewy neurite-like aggregates. Nat Protoc 2014, 9(9):2135–2146.

26. Beaudoin GM, 3rd, Lee SH, Singh D, Yuan Y, Ng YG, Reichardt LF, Arikkath J: Culturing pyramidal neurons from the early postnatal mouse hippocampus and cortex. Nat Protoc 2012, 7(9):1741–1754.

27. Giasson BI, Duda JE, Quinn SM, Zhang B, Trojanowski JQ, Lee VMY: Neuronal α-Synucleinopathy with Severe Movement Disorder in Mice Expressing A53T Human α-Synuclein. Neuron 2002, 34(4):521–533.

28. Kuo YM, Li Z, Jiao Y, Gaborit N, Pani AK, Orrison BM, Bruneau BG, Giasson BI, Smeyne RJ, Gershon MD et al: Extensive enteric nervous system abnormalities in mice transgenic for artificial chromosomes containing Parkinson disease-associated alpha-synuclein gene mutations precede central nervous system changes. Hum Mol Genet 2010, 19(9):1633–1650.

29. Williams JM, Duckworth CA, Vowell K, Burkitt MD, Pritchard DM: Intestinal Preparation Techniques for Histological Analysis in the Mouse. Curr Protoc Mouse Biol 2016, 6(2):148–168.

30. Dunn KW, Kamocka MM, McDonald JH: A practical guide to evaluating colocalization in biological microscopy. Am J Physiol Cell Physiol 2011, 300(4):C723–742.

31. Santos J, Gracia P, Navarro S, Peña-Díaz S, Pujols J, Cremades N, Pallarès I, Ventura S: α-Helical peptidic scaffolds to target α-synuclein toxic species with nanomolar affinity. Nature Communications 2021, 12(1):3752.

32. Mikula H, Stapleton S, Kohler RH, Vinegoni C, Weissleder R: Design and Development of Fluorescent Vemurafenib Analogs for In Vivo Imaging. Theranostics 2017, 7(5):1257-1265.

33. Demétrio de Souza França P, Kossatz S, Brand C, Karassawa Zanoni D, Roberts S, Guru N, Adilbay D, Mauguen A, Valero Mayor C, Weber WA et al: A phase I study of a PARP1-targeted topical fluorophore for the detection of oral cancer. Eur J Nucl Med Mol Imaging 2021, 48(11):3618-3630.

34. Ehrnhoefer DE, Bieschke J, Boeddrich A, Herbst M, Masino L, Lurz R, Engemann S, Pastore A, Wanker EE: EGCG redirects amyloidogenic polypeptides into unstructured, off-pathway oligomers. Nature Structural & Molecular Biology 2008, 15(6):558–566.

35. Chen SW, Drakulic S, Deas E, Ouberai M, Aprile FA, Arranz R, Ness S, Roodveldt C, Guilliams T, De-Genst EJ et al: Structural characterization of toxic oligomers that are kinetically trapped during α-synuclein fibril formation. Proceedings of the National Academy of Science 2015, 112:E1994–E2003.

36. Polinski NK, Volpicelli-Daley LA, Sortwell CE, Luk KC, Cremades N, Gottler LM, Froula J, Duffy MF, Lee VMY, Martinez TN et al: Best Practices for Generating and Using Alpha-Synuclein Pre-Formed Fibrils to Model Parkinson’s Disease in Rodents. J Parkinsons Dis 2018, 8(2):303–322.

37. Maillard J, Klehs K, Rumble C, Vauthey E, Heilemann M, Fürstenberg A: Universal quenching of common fluorescent probes by water and alcohols. Chem Sci 2020, 12(4):1352–1362.

38. Ojida A, Sakamoto T, Inoue M-a, Fujishima S-h, Lippens G, Hamachi I: Fluorescent BODIPY-Based Zn(II) Complex as a Molecular Probe for Selective Detection of Neurofibrillary Tangles in the Brains of Alzheimer’s Disease Patients. Journal of the American Chemical Society 2009, 131(18):6543–6548.

39. van der Sommen F, Curvers WL, Nagengast WB: Novel Developments in Endoscopic Mucosal Imaging. Gastroenterology 2018, 154(7):1876–1886.

40. Thakur P, Breger LS, Lundblad M, Wan OW, Mattsson B, Luk KC, Lee VMY, Trojanowski JQ, Björklund A: Modeling Parkinson’s disease pathology by combination of fibril seeds and α-synuclein overexpression in the rat brain. Proceedings of the National Academy of Sciences 2017, 114(39):E8284–E8293.

41. Rota L, Pellegrini C, Benvenuti L, Antonioli L, Fornai M, Blandizzi C, Cattaneo A, Colla E: Constipation, deficit in colon contractions and alpha-synuclein inclusions within the colon precede motor abnormalities and neurodegeneration in the central nervous system in a mouse model of alpha-synucleinopathy. Translational Neurodegeneration 2019, 8(1):5.

42. Majbour NK, Vaikath NN, Eusebi P, Chiasserini D, Ardah M, Varghese S, Haque ME, Tokuda T, Auinger P, Calabresi P et al: Longitudinal changes in CSF alpha-synuclein species reflect Parkinson’s disease progression. Movement Disorders 2016, 31(10):1535–1542.

43. Stewart T, Sossi V, Aasly JO, Wszolek ZK, Uitti RJ, Hasegawa K, Yokoyama T, Zabetian CP, Leverenz JB, Stoessl AJ et al: Phosphorylated α-synuclein in Parkinson’s disease: correlation depends on disease severity. Acta Neuropathologica Communications 2015, 3(1):7.

44. Siderowf A, Concha-Marambio L, Lafontant D-E, Farris CM, Ma Y, Urenia PA, Nguyen H, Alcalay RN, Chahine LM, Foroud T et al: Assessment of heterogeneity among participants in the Parkinson’s Progression Markers Initiative cohort using alpha-synuclein seed amplification: a cross-sectional study. The Lancet Neurology 2023, 22(5):407–417.

45. Russo MJ, Orru CD, Concha-Marambio L, Giaisi S, Groveman BR, Farris CM, Holguin B, Hughson AG, LaFontant D-E, Caspell-Garcia C et al: High diagnostic performance of independent alpha-synuclein seed amplification assays for detection of early Parkinson’s disease. Acta Neuropathologica Communications 2021, 9(1):179.

46. Roos DS, Klein M, Deeg DJH, Doty RL, Berendse HW: Prevalence of Prodromal Symptoms of Parkinson’s Disease in the Late Middle-Aged Population. J Parkinsons Dis 2022, 12(3):967–974.

47. Chan PW, Ngu JH, Poh Z, Soetikno R: Colorectal cancer screening. Singapore Med J 2017, 58(1):24–28.

48. Shi J, Wang Y, Chen D, Xu X, Li W, Li K, He J, Su W, Luo Q: The alteration of intestinal mucosal α-synuclein expression and mucosal microbiota in Parkinson’s disease. Applied Microbiology and Biotechnology 2023, 107(5):1917–1929.

