## Supporting Information for "Fluorescent Peptide-based Probe for the Detection of Alpha-synuclein Aggregates in the Gut"

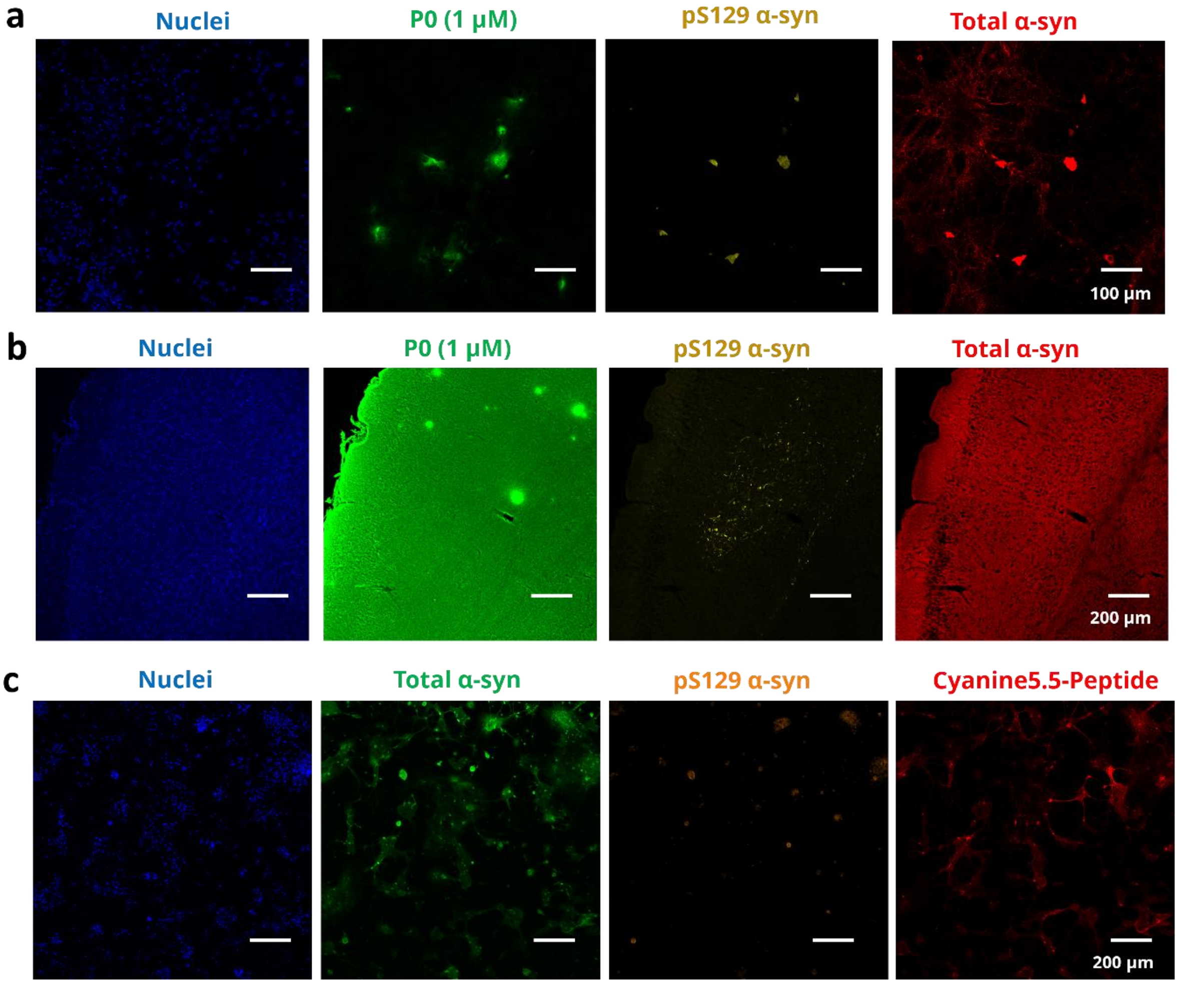


Supplementary Figure 1. *In vitro* and *ex vivo* staining of peptide-based fluorescent probe **P0** and cyanine5.5 labelled peptide. (a) Confocal images showing staining of probe **P0** (green) in comparison with phosphorylated α-syn antibody staining (yellow) in primary hippocampal neuronal cells treated with PFFs; (b) Confocal images showing staining of probe **P0** in brain tissues with high tissue autofluorescence in the green channel making it difficult to visualize specific staining; (c) Confocal images showing staining of cyanine5.5-labelled peptide (red) showing non-specific binding in primary hippocampal neuronal cells, as it binds to α-syn monomers not labelled by pS129 antibody.

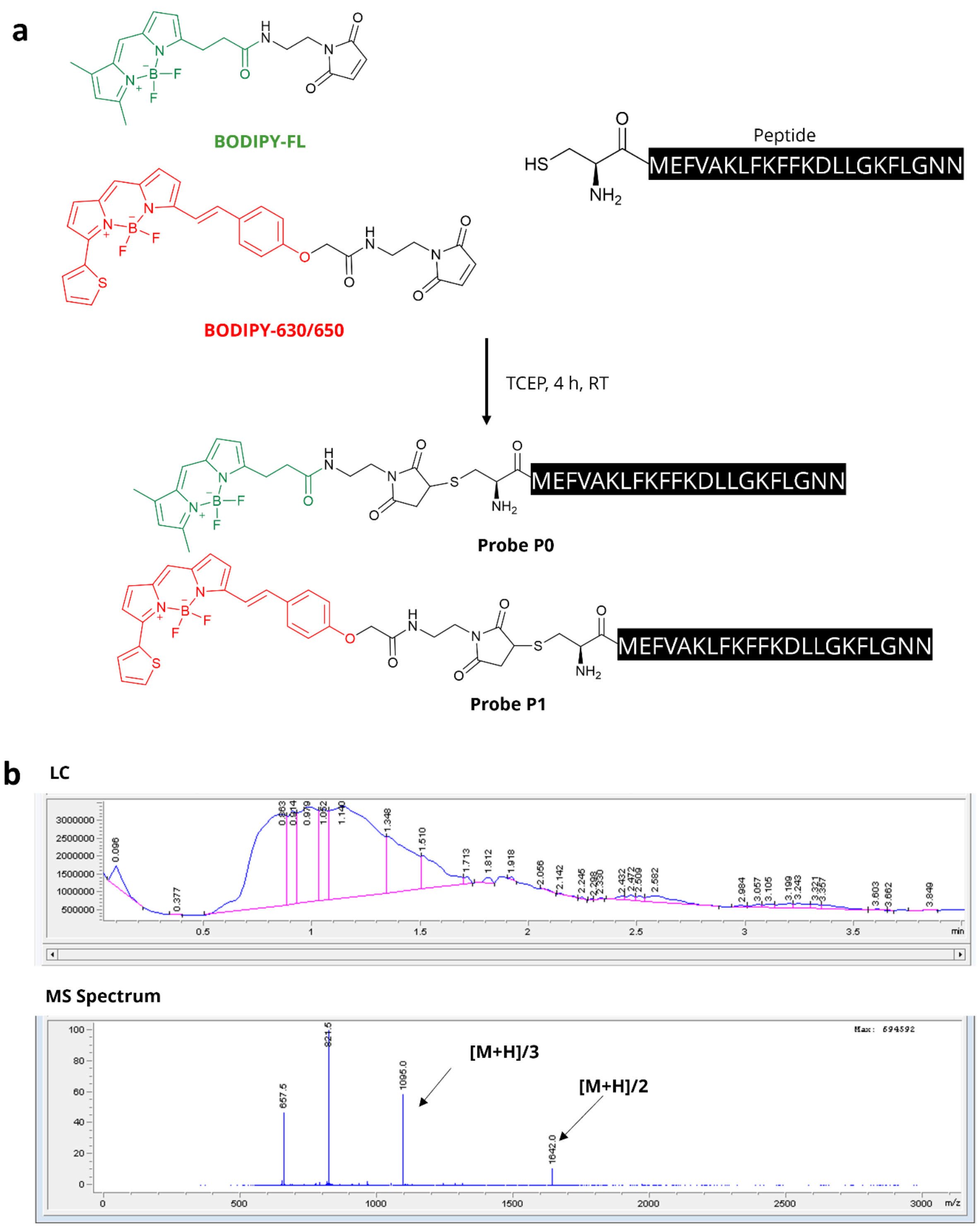


Supplementary Figure 2. Synthesis and characterization of peptide-based fluorescent probes **P0** and **P1**. (a) Synthesis of probes using BODIPY-dye (BODIPY-FL maleimide for **P0**, BODIPY630/650 for **P1**), TCEP, peptide in water, reaction for 4 h at room temperature; (b) LCMS characterization of purified probe **P1**. Single peak in LC shows purified P1 and corresponding MS spectrum shows [M+H]/2 peak of 1642.0 Da and [M+H]/3 peak of 1095.0 Da, according to the expected molecular weight of P1, 3282.68 Da. [M+H] not seen in the spectrum as the maximum mass that can be measured on the spectrometer is 3000 Da.

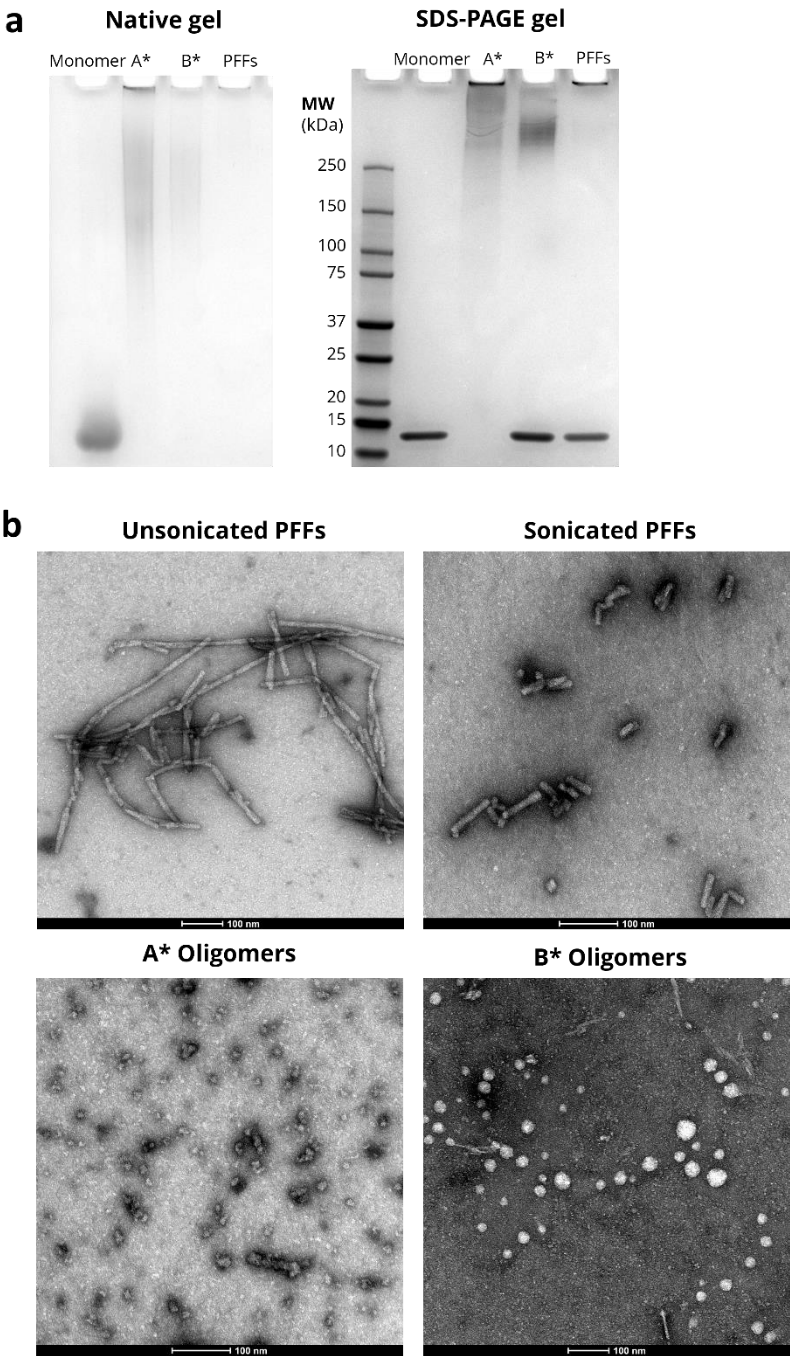


Supplementary Figure 3. Characterisation of various α-syn aggregates. (a) Native-PAGE and (b) SDS-PAGE gel showing α-syn monomers, A* oligomers, B* oligomers and PFFs; (b) TEM images showing the size and morphology of the aggregates. The unsonicated PFFs are large (>200 nm), while sonicated PFFs are around 50-60 nm. Spherical structures (10-20 nm) are observed for the A* oligomers. Slightly larger (20-30 nm) spherical structures are observed for the B* oligomers.

a

b

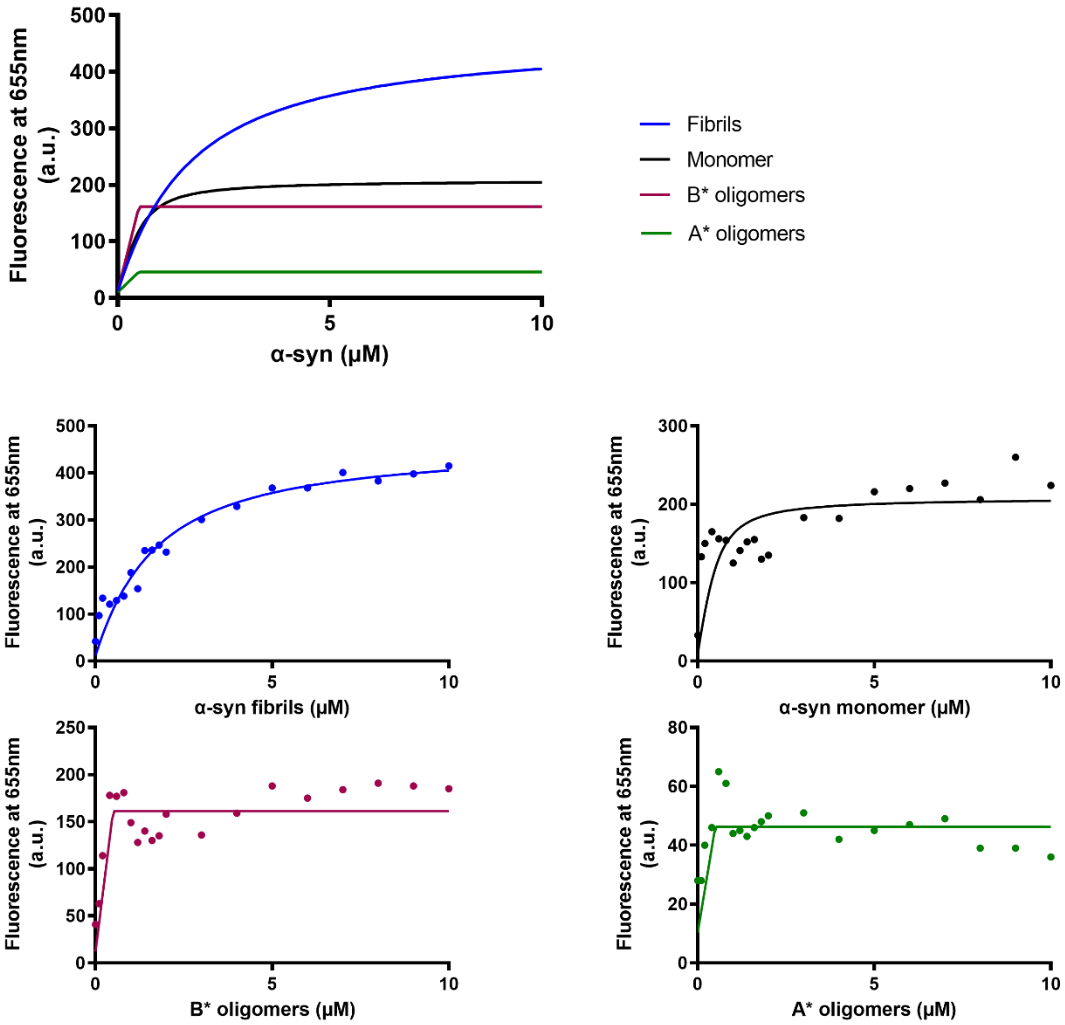


c

| α-syn species | $K_D$ (μM) | $R^2$ |
|---|---|---|
| **Fibrils (PFFs)** | 1.414 ± 0.2581 | 0.9397 |
| **Monomers** | N.D. | 0.2460 |
| **B* oligomers** | N.D. | 0.5836 |
| **A* oligomers** | N.D | -0.03741 |

Supplementary Figure 4. (a) Combined fluorescence titration curve with various α-syn species (monomers, A* and B* oligomers); (b) Individual titration curves of each α-syn species. (c) Table of $K_D$ and $R^2$ values of each curve. Only α-syn fibrils produce a physically relevant and reliable $K_D$ and $R^2$ readout. $K_D$ is not determined (N.D.) for monomers, A* and B* oligomers as the $R^2$ value of the curves indicated that the curves were not well fitted. As is common in literature, it is challenging to design a fluorescent probe that demonstrates sufficient fluorescent enhancement in the presence of α-syn oligomers. Our current fluorescent assays are insufficiently sensitive to detect probe-oligomer binding.

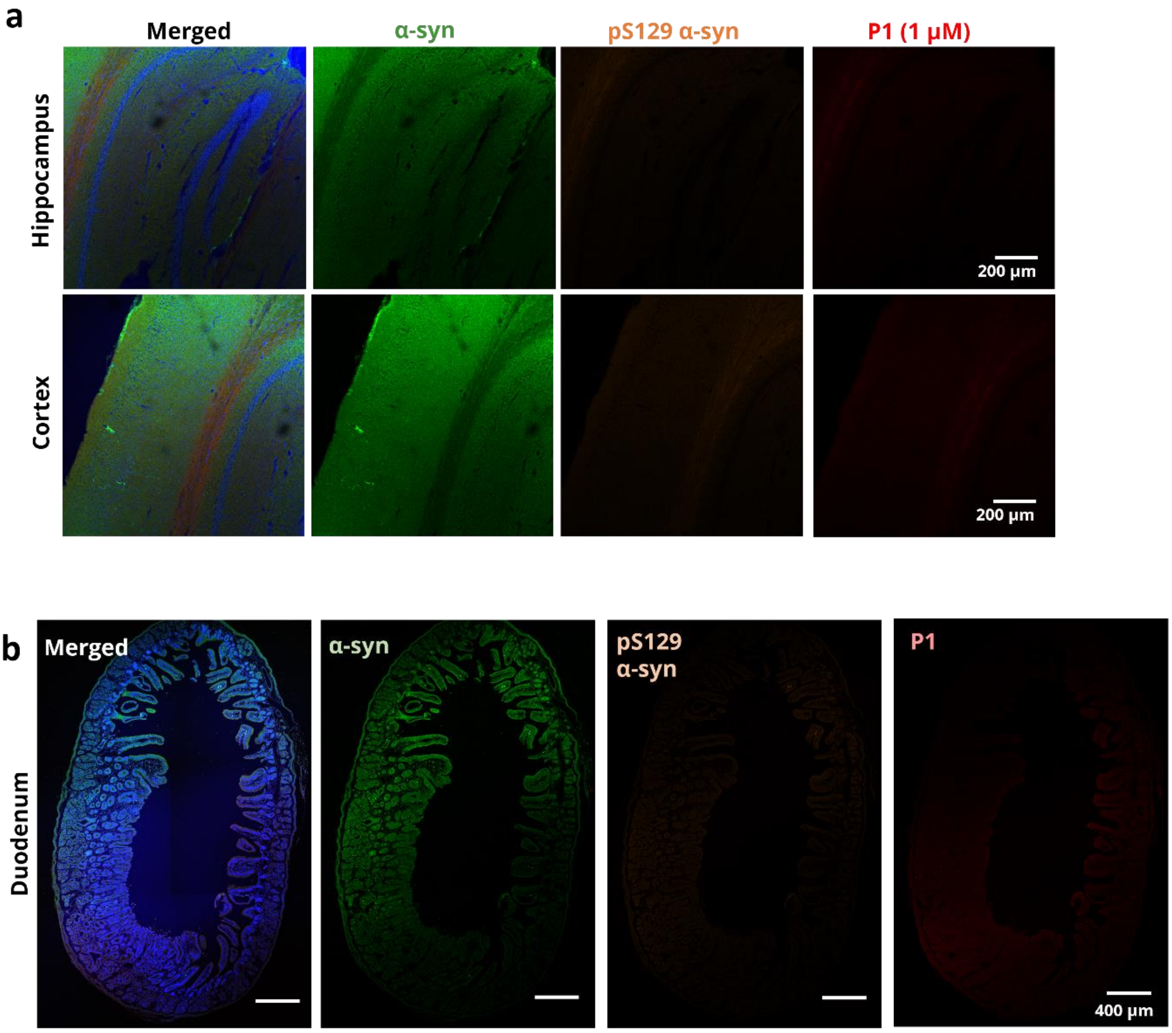


Supplementary Figure 5. (a) Confocal microscopy of brain tissues post 14 days of PBS injection showing no detectable levels of α-syn aggregation in the cortex region; (b) Confocal microscopy of duodenum tissues post 75 days of PBS injection showing low levels of α-syn aggregation in the mucosa region indicated by the white arrows. Nuclei in blue, α-syn in green, pS129 α-syn in orange and probe in red.

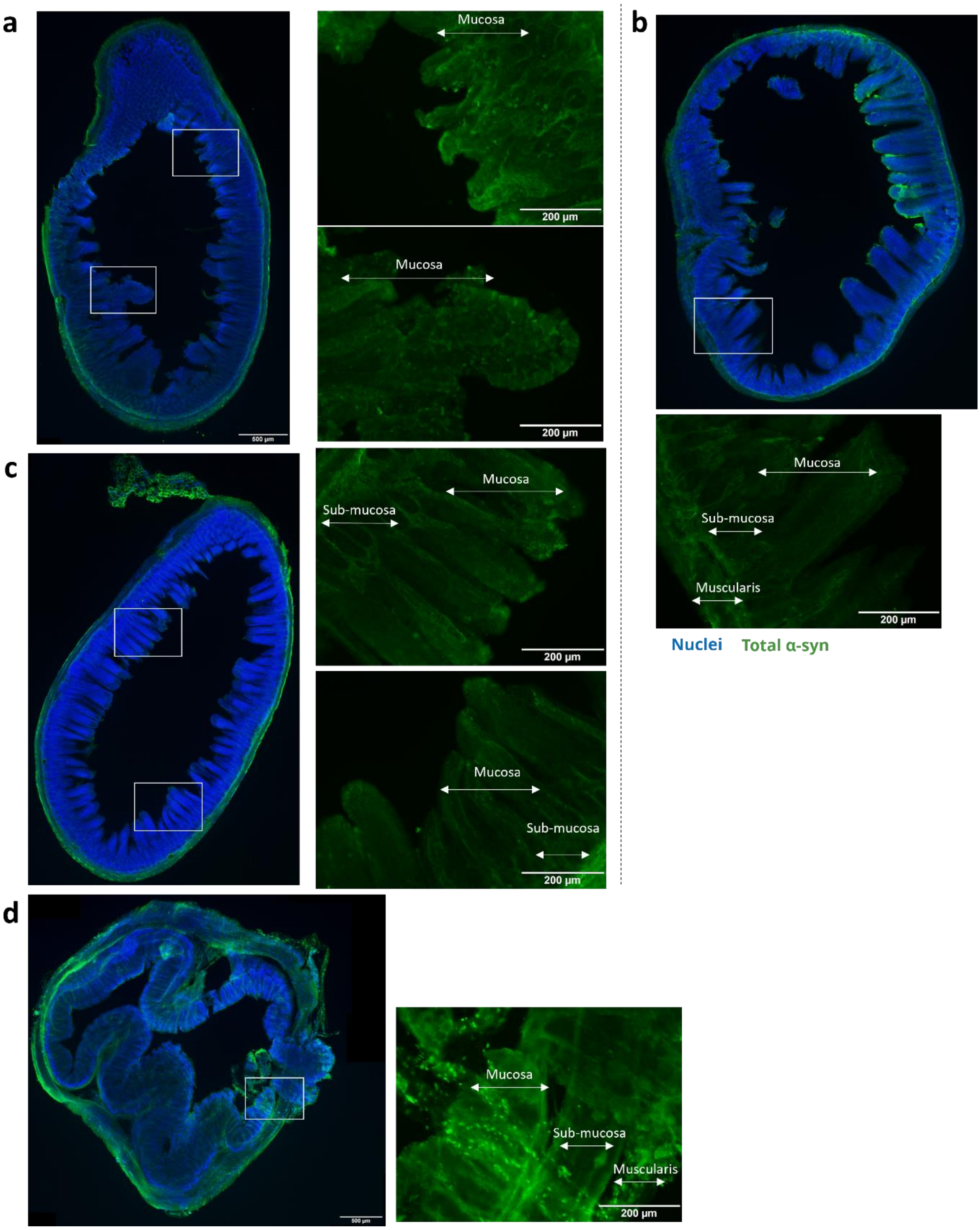


Supplementary Figure 6. Immunofluorescence staining showing the presence of α-syn expression throughout the GI tract and across the tissue layers (muscularis externa, submucosa, mucosa) in wild-type mouse (a) duodenum, (b) jejunum, (c) ileum and (d) colon.

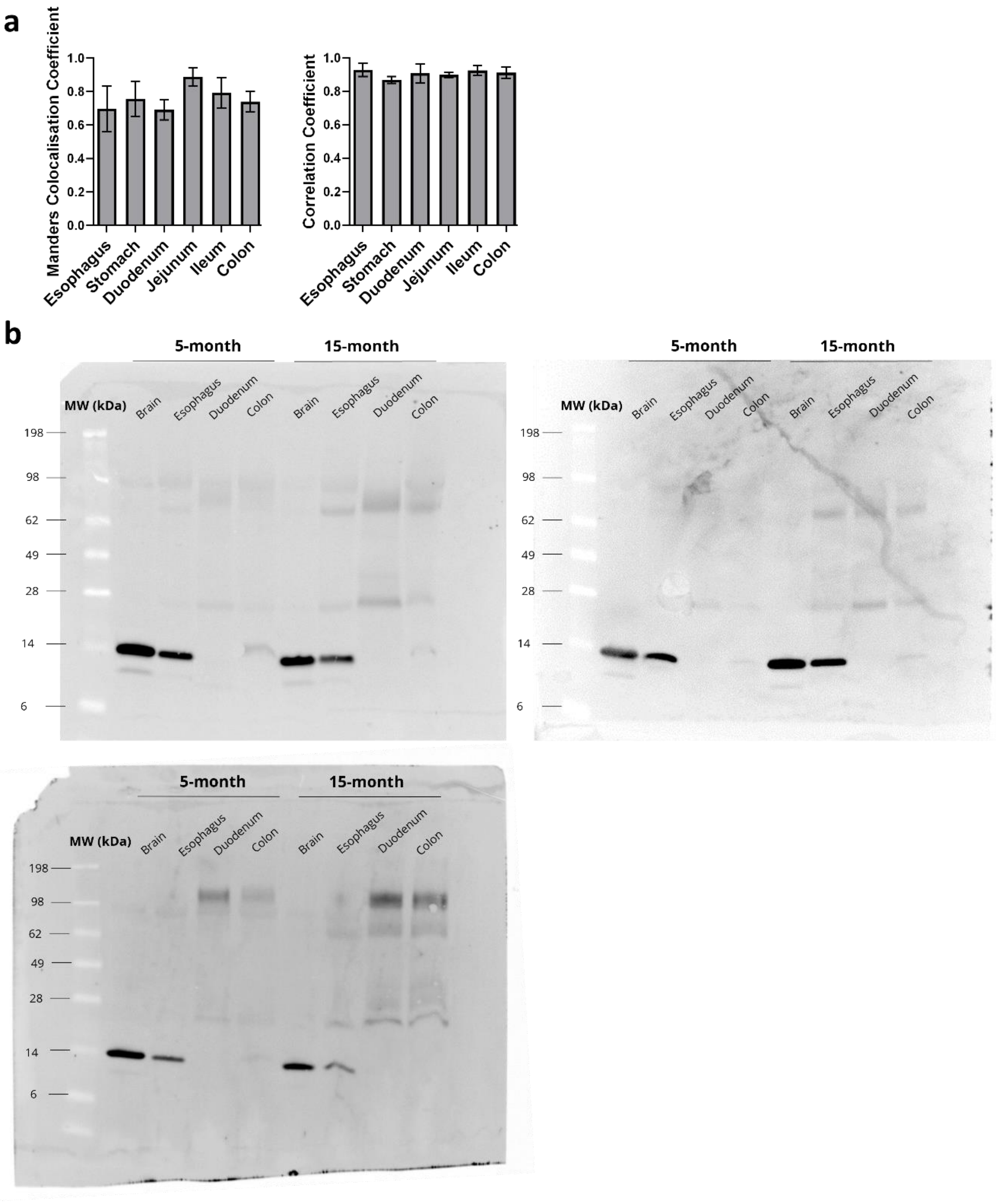


Supplementary Figure 7. (a) Graphs showing the Manders colocalization and Pearson's correlation coefficients calculated between pS129 α-syn antibody and probe **P1** channels; (b) Full, uncropped western blots (3 replicates) of protein lysates from tissues of transgenic mice.

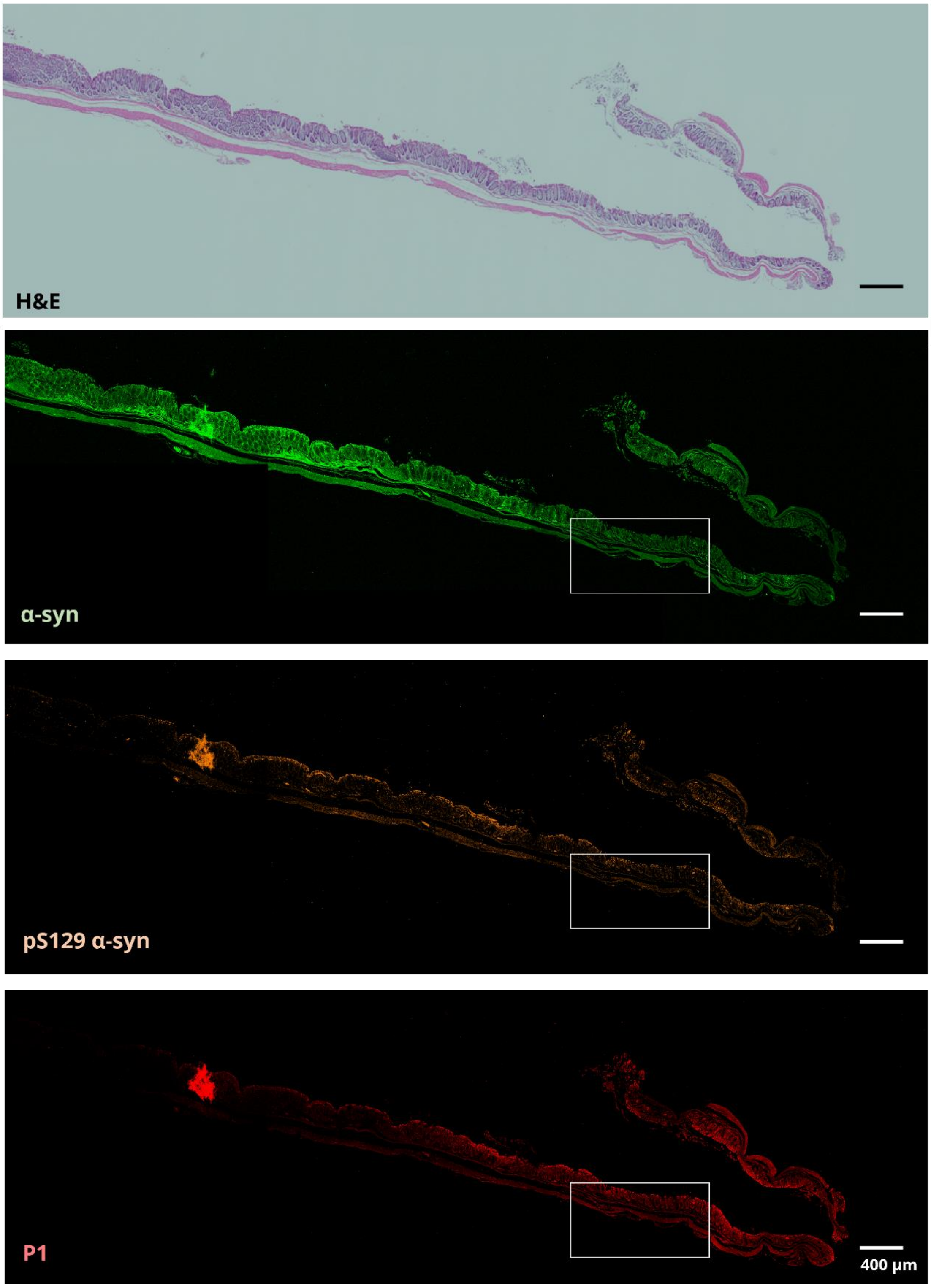


Supplementary Figure 8. Longitudinal section of transgenic mouse colon. H&E and fluorescence images showing the entire length of colon tissue used for quantitative analysis, with the white boxes indicating the region that is highlighted in Fig 6.
